# Deep tissue removal in wounds facilitates algal colonization and inhibits healing and regeneration in tropical corals

**DOI:** 10.64898/2026.07.16.738947

**Authors:** Ashley W. Seifert, Molly Brzezinski, Craig W. Osenberg, Adrian C. Stier

**Affiliations:** Department of Biology, University of Kentucky, Lexington, KY 40508; Department of Ecology, Evolution, and Marine Biology, University of California Santa Barbara, Santa Barbara, California 93106, USA; Marine Science Institute, University of California Santa Barbara, Santa Barbara, California 93106, USA; Odum School of Ecology, University of Georgia, Athens, GA, USA

## Abstract

Although corals are highly regenerative, some colonies in reef ecosystems completely recover from sublethal damage while other colonies exhibit partial mortality to similar injuries. To understand factors that might naturally curtail regenerative ability, we experimentally wounded small colonies in three coral genera (*Acropora*, *Pocillopora*, *Porites*) by mimicking natural corallivory using scraping (tissue and skeletal damage) or airbrushing (deep tissue removal with no skeletal injury). We found all scraped wounds regenerated rapidly in *Acropora* and *Porites*, while *Pocillopora* fragments frequently retained open lesions. In stark contrast, airbrushing resulted in algal colonization and delayed tissue healing and regeneration across all corals. Detailed cellular analysis of *Porites* wounds revealed two general phases comprising tissue regeneration: a healing phase defined by rapid coverage of bare skeleton with coenosarc, pigment cells and gastrodermal reformation, and then a second phase lasting one week ending in polyp regeneration. Red fluorescence appeared transiently in scrape wounds but persisted in tissue at the wound margin surrounding algae in airbrush wounds, suggesting that algal occupation of the wound bed inhibits coenosarc healing. Lastly, histological cross-sections of healing airbrush wounds in *Porites* revealed progressive loss of deep tissue leading to skeletal breakdown beneath the wound. Together, our results demonstrate the biphasic nature of tissue regeneration in colonial corals and provide a framework for understanding how biotic factors impact tissue repair and regeneration in nature.

## INTRODUCTION

Generally speaking, cnidarians exhibit incredible regenerative ability in response to tissue damage (Bosch 2007; Luz et al. 2021; Han et al. 2025) and free-living cnidarians like *Hydra* (Hydrozoa) and *Nematostella* (Anthozoa) along with the colonial cnidarian *Hydractinia* (Hydrozoa) have been a well-spring of knowledge for understanding how stem cells support homeostatic tissue maintenance and reparative regeneration (Bosch 2007) (Figure 1A-B). Specifically, *Hydractinia* mobilize pluripotent stem cells, while *Hydra* and *Nematostella* direct lineage-restricted, multipotent stem cells to replace missing tissue with all three models appearing to show negligible declines in regenerative ability after repeated injury. Cnidarians, however, are far more diverse than these model species and the ideal laboratory conditions under which they are studied make it difficult to tease apart how tissue repair mechanisms may vary under more natural conditions. For example, stony corals (Scleractinia) are the foundation of marine reef ecosystems where they are susceptible to a range of injuries and yet their natural regenerative ability remains curiously understudied at the cellular and molecular level. Corals frequently sustain sublethal injuries from predation, physical disturbance, and disease (Loya 1976; Bak and Criens 1982; Bythell et al. 1993; Kramarsky-Winter and Loya 2000) and while severity may vary, all injuries cause tissue damage or loss.

**Figure 1.**
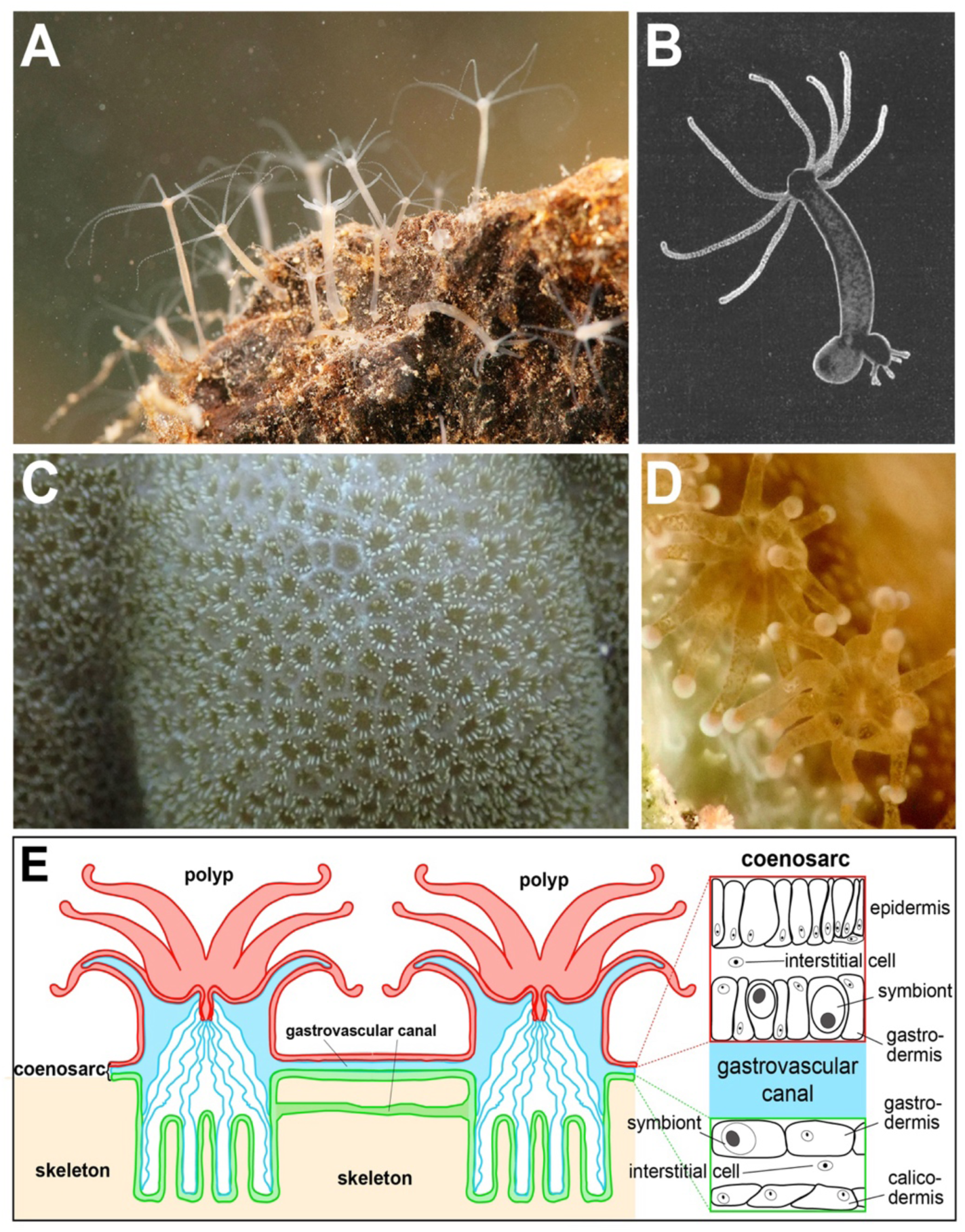
Comparison of single polyp and multi-polyp cnidarians. Comparative anatomy and organization of cnidarian body plans relevant to coral wound healing. **(A)** Collection of individual *Hydra* polyps affixed to rock substrate. **(B)** Single, free-living *Hydra* polyp showing columnar organization with tentacles and mouth at the anterior (top) and foot at the posterior (bottom). An asexual bud is visible emerging from the posterior. **(C)** Surface view of a perforate stony coral (*Porites*) colony showing individual polyps residing within regularly spaced skeletal cups. **(D)** High-magnification image of two adjacent *Porites* polyps extending tentacles outside the skeleton during feeding or defense. **(E)** Diagram of typical *Porites* colony organization, showing polyps joined by interstitial tissue (coenosarc) comprised of an outer (red) layer of tissue exposed to seawater and an inner (green) cellular bi-layer of dermis resting atop the calcium-carbonate skeleton (beige). The space between these tissue layers forms the gastrovascular cavity and gastrovascular canals between polyps (blue). Insets provide cellular organization of the coenosarc including epidermis, gastrodermis, interstitial cells and symbiotic dinoflagellates. *Panel E adapted from information provided by the Coral Disease and Health Consortium (CDHC), Murphy McDonald, NOAA*.

Most stony corals are colonial, exhibit a mutualistic symbiosis with microscopic algae, and secrete a calcium carbonate skeleton; traits that differentiate them from the non-calcifying cnidarian models mentioned above. Although most stony corals possess algal symbionts, recent phylogenetic analysis supports that the common ancestor of all cnidarian lineages was a free-living polyp that lacked symbionts (Vaga et al. 2025) and evidence supports that symbionts are not required for tissue regeneration (DeFilippo et al. 2016; Burmester et al. 2017). In stony corals, polyps are nested inside individual skeletal cups (calyxes) with deeper skeleton beneath and are linked by interstitial tissue (coenosarc) of varying complexity (Figure 1C-E). In imperforate corals, the coenosarc exists as a double bilayer of tissue anchored atop a dense skeleton. In perforate corals, a thicker (∼5mm) interconnected web of gastrodermal and mesenterial tissue extends through a porous skeleton and supports superficial polyps. In some perforate corals, an endolithic layer of algae exists just below the living coral tissue (del Campo et al. 2017). Although these endolithic algae may support coral recovery by modulating skeletal reflectance after bleaching stress (Galindo-Martínez et al. 2022), the possibility of endolithic algal blooms to impede healing in deep wounds has yet to be tested directly.

Skeletal architecture and corresponding tissue interconnectedness may lead to substantial disparity in how wound healing and regeneration proceeds in colonial stony corals compared to single-polyp or non-calcifying cnidarians. Wound architecture affects the kinetics of tissue migration and the rate of healing, which in turn can affect colony survival and fitness. For example, some corallivores inflict injuries that remove small bits of tissue without damaging the skeleton. Gastropods (e.g., *Drupella* spp., *Coralliophila* spp.) consume tissue while leaving the skeleton intact, and crown-of-thorns seastars (*Acanthaster planci*) remove large amounts of tissue without damaging the skeleton. In contrast, parrotfishes (Scaridae) and pufferfishes (Tetraodontidae) abrade or excavate tissue and cause skeletal damage (Cameron and Edmunds 2014; Palacios et al. 2014), not unlike damage caused by physical disturbances like sand abrasion or storm-induced fragmentation. Wounds that heal more slowly are more likely to be colonized by fouling organisms such as algae and vermetids that settle onto the exposed skeleton (Phillips et al. 2014). Once established, fouling organisms can prevent or slow tissue regrowth, possibly resulting in permanent, localized tissue mortality (Bak and Steward-Van Es 1980; Meesters et al. 1997; Kramarsky-Winter and Loya 2000; Titlyanov et al. 2005). Thus, understanding how the extent of tissue loss, with or without skeletal damage, affects healing and regeneration requires further study within and across species.

Understanding how corals with different tissue and skeletal architectures heal under different wound damage is critical given the escalating biotic and abiotic stressors being imposed on coral reefs (Kramarsky-Winter and Loya 2000; Palmer et al. 2011), and the central role that corals play in these valuable ecosystems. Previous studies comparing rates of healing among different wound types have yielded conflicting results. In one study, tissue only lesions healed faster than scrapes which removed tissue and skeleton in *Agaricia agaricites* and *Porites astreoides*, although in *Porites* skeletal damage initially accelerated regeneration (Bak and Steward-Van Es 1980). Similar results were obtained by Cameron and Edmunds (2014) who found that shallow tissue removal wounds healed faster than those resulting from tissue removal and skeletal abrasion, although this effect was confounded by the size of the wounds (i.e., tissue-only wounds were ∼⅓ the size of the tissue plus skeletal removals). In contrast, other studies found species responded differently to tissue removal versus scrapes: *Porites australiensis* healed faster when the skeleton was scraped, while *Acropora hyacinthus* and *Montipora tuberculosa* showed the opposite pattern, and *Acropora cytherea* showed no difference (Hall 1997, 2001). Furthermore, Hall (2001) found that regeneration rate and algal colonization were negatively correlated, suggesting that slower regeneration may lead to high rates of partial or complete mortality due to the deleterious effects of algae. Hall (1998, 2001) and Bak & Steward-Van Es (1980) collectively established that wound type and species architecture both shape coral healing, but their designs confounded wound size and depth. No study has experimentally separated tissue-removal depth from skeletal damage to investigate what drives algal colonization of the wound bed — we do that here.

Incorporating fixed wound sizes and types in our studies (see below), we examine three hypotheses: (1) Regenerative ability varies between species with different skeletal architecture such that perforate corals with access to deep tissue beneath wounds will heal and regenerate more rapidly from surface scrapes because residual tissue within the wound bed can rapidly repopulate the wound bed and facilitate regeneration. In contrast, imperforate corals will be dependent on lateral tissue invasion and will heal more slowly (Bak and Steward-Van Es 1980; Hall 1997, 2001) (2) Deep tissue removal in perforate corals is more vulnerable to colonization by turf algae because lateral wound healing takes longer compared to wounds that can also be repopulated by the growth of remnant tissue in the wound bed. (Meesters et al. 1997; Hall 2001; McCook et al. 2001; Titlyanov et al. 2005). (3) Skeletal damage associated with wounding will affect healing and regeneration (negatively or positively) compared to similar wound types where the skeleton is not damaged. Although an exact mechanism has not been demonstrated, skeletal damage is associated with short-term accelerated growth and high calcification rates at the injury margin in some species and these processes may facilitate faster tissue regeneration (Lock et al. 2022). Alternatively, because skeletal microdamage can locally increase pH, a more alkaline local environment will support growth of pathogenic microbes and algae which would reduce the healing rate. These hypotheses are not independent from one another but differ to some extent in their assertions about the role of skeletal damage vs. tissue removal.

With these hypotheses in mind, we evaluated tissue regeneration in three common Indo-Pacific coral genera in response to two injury types that mimic distinct modes of damage by corallivores. To evaluate coral tissue regeneration, we measured patterns of coenosarc regrowth and polyp regeneration, quantified algal colonization, and characterized cellular dynamics of wound healing. Throughout our work, we make the distinction between wound healing and tissue regeneration. We use the term “healing” to describe the reestablishment of the epithelial barrier in the wound bed, which is first demonstrated as the appearance of coenosarc covering the wound bed. By contrast, “regeneration” is used to describe the complete replacement of polyps and coenosarc within the wound bed.

## METHODS

### Site and Species Description

We carried out studies at the Gump Research Station in Mo’orea, French Polynesia. We focused on three coral taxa: *Porites* spp, *Acropora hyacinthus*, and *Pocillopora* spp. *Acropora hyacinthus* is a fast-growing, branching coral with a fragile, imperforate skeleton and thin tissue. *Pocillopora* are branching species with a dense skeletal framework and moderately thick tissue. *Porites* species are slow-growing massive corals with thick tissue and a perforate skeleton. All three genera comprise cryptic or morphologically convergent species lineages that we could not reliably distinguish in the field — *Pocillopora* (Johnston et al. 2022), the *Acropora hyacinthus* complex (Rassmussen et al. 2025), and massive *Porites* (Primov et al. 2024) — so we refer to all three by genus throughout. All Porites colonies used in Experiments 2 and 3 were of the massive growth form characteristic of *P. lobata*/*P. lutea* morphospecies; we did not include any encrusting morphotypes (e.g., *P. rus*), and we did not collect tissue samples for molecular species-level confirmation.

### General Experimental Details

In our experiments, we used two basic wound types that mimic injuries created by different predators that have been contrasted in prior studies: (1) scraping wounds (produced with a Dremel tool) that damaged skeleton and tissue, which simulate parrotfish grazing; and (2) deep tissue wounds (produced with an airbrush) that removed tissue more deeply but left the skeleton intact. These airbrush wounds mimic invertebrate predation by snails and sea stars. We refer to these wound types as “scraped” and “airbrushed”.

The first set of experiments (Experiment 1) investigated the extent of regeneration and algal colonization in three coral species across the two wound types (scraped and airbrushed). The second set of experiments (Experiment 2) focused on *Porites* (a perforate coral species where coral tissue is connected via deep channels in the skeleton and on the surface - see Figure 1E). Experiment 2 was designed to assess the relative role of total tissue removal vs. skeletal abrasion to evaluate competing hypotheses to explain the faster regeneration commonly observed when wounds are created by scraping. Specifically, we imposed three wounding treatments: airbrushing, scraping, and a combination wound made by airbrushing followed by scraping. The performance of corals in the third treatment, relative to the other two, allowed us to evaluate if the faster regeneration we observed in scraped wounds was due to skeletal damage. As in Experiment 1, we followed the healing process using microscopy and monitored algal colonization. Additionally, we assessed patterns of red/green fluorescence in healing tissue and at wound margins. In our final study (Experiment 3), we conducted a wounding experiment using *Porites* fragments where we made either scrape or airbrush wounds (as in Experiments 1 & 2) and then fixed the entire fragment at specific time points for cellular analysis using standard histological techniques to explore cell migration and deeper tissue at the margins of the wound bed.

### Experiment 1: Effect of wound type on regeneration across three coral species

#### Collection and Wounding

We collected two fragments each from 11 different parent colonies (5 *Acropora*, 3 *Pocillopora*, and 3 *Porites* colonies), transported them to the Gump Research Station, and acclimated them in shaded, flow-through seawater tables for 48 hours before experimental manipulation. One fragment from each colony was assigned to one of the two treatments (i.e., wound types). Scrape wounds were created by abrading approximately the top 1 mm of surface tissue and skeleton with a handheld Dremel equipped with a fine diamond burr tip. Deeper tissue-only wounds were created using an airbrush. Wound size was standardized across treatments using a 1 × 1 cm silicone stencil placed centrally on the upper surface of each fragment. The experiment was conducted in June 2022, during the austral winter.

#### Response Variables

Following injury, we photographed each wound under bright field illumination at 0.63× and 1.25× magnification on day 0 (immediately after wounding, D0), at 72 hours (D3), and every five days thereafter until day 28 (D8 - D28). We evaluated wound regeneration by classifying the wound as completely regenerated (all portions of the wound were covered by well-pigmented coral tissue and mature polyps). At every imaging timepoint we also scored algal colonization of each wound as a binary state: present (if filamentous or turf algae occupied the wound bed) or absent. We did not quantify algal cover or identify algal taxa and applied the same binary scoring in Experiments 2 and 3.

#### Analysis

We modelled two outcomes from Experiment 1: Complete tissue regeneration (the reappearance of polyps within the wound area) and algal colonization of the wound bed, each with a binomial GLMM of wound type, species, and their interaction. The wound type × species interaction was negligible for regeneration (likelihood-ratio test: χ² = 2.07, df = 2, P = 0.355) but significant for algal colonization (χ² = 9.28, df = 2, P = 0.0096); because the wound-type effect on algae was nonetheless large and consistent across genera, we report additive-model effects for both. Because no *Pocillopora* scrape wounds regenerated, a standard logistic model could not estimate these odds ratios; we therefore estimated wound-type and species odds ratios for both outcomes with Firth’s penalized logistic regression (Kosmidis and Firth 2021), which gives finite estimates when one group is all-or-nothing, and report it as the primary model; a standard mixed model gave the same direction and magnitude. For complete coenosarc healing, we report the standard binomial mixed model as the primary model for that outcome and Firth’s penalized regression as a robustness check. We obtained model-predicted probabilities and within-species differences between wound types with estimated marginal means (Lenth 2023). We confirmed that these conclusions held across six missing-data scenarios and after excluding one fragment with an irregular starting wound. A leave-one-out refit excluding a single coral confirmed that the wound-type effect is robust, although the genus difference in complete healing depended on this fragment. All Experiment 1 analyses used R version 4.5.2. Inference was based on the binomial GLMM fitted to repeated wound-state assessments across all post-injury timepoints (days 0, 3, 8, 13, 18, 23, 28) with fragment identity as a random intercept; we did not model time as a fixed effect but instead analyzed the endpoint (Tables S1–S9; Figure S3). A day-28 endpoint Fisher exact test corroborated the wound-type effect model-free (within Acropora, five of five scrape versus zero of five airbrush fragments healed, P = 0.008).

### Experiment 2: Separating effects of skeletal damage vs deep tissue removal on healing and regeneration

#### Collection and Wounding

Fifteen juvenile colonies of *Porites* were collected along the northwest shore of Mo’orea during the austral winter of 2025. Colonies were gently chiseled off pavement and transported back to Gump station where their bases were leveled with a bandsaw (Gryphon C-40 Aquasaw) and glued to labeled tiles. Corals were acclimated in flow-through seawater tables for one day at stable conditions (27-28°C temperature, salinity ∼35 PSU) before experimental manipulation. Corals were wounded a single time using scraping or airbrushing as in Experiment 1, with wound size standardized to 8 mm diameter circles using a silicone stencil as in Experiment 1. Combination (airbrush + scrape) wounds were created by removing tissue with an airbrush and then damaging skeleton with the Dremel over the same airbrush wound. Following wounding, we maintained coral fragments in shaded, flow-through seawater tables under ambient photoperiod conditions (∼11-hour daylight cycles).

#### Response Variables

To capture detailed early-stage healing responses, each colony was imaged using an SZX-10 Olympus stereomicroscope equipped with filters for fluorescence (GFP- 470nm and RFP- 545nm) and a high-resolution camera. Corals were imaged immediately after wounding (D0), daily for five days (D1 through D5), and at weekly intervals until D63 (starting at D7) or until completely regenerated. We conducted imaging at two standardized magnifications: one (1.25×) to characterize the coverage by coenosarc and polyps and another at higher magnification (3.2×) to more precisely observe polyp emergence and fluorescence signals at wound margins. Brightfield and fluorescent imaging was performed under moderate darkness (Gain = 1, exposure = 100-300ms, LED fluorescent light intensity = 50%). Corals were imaged in seawater and immediately returned to the seawater table.

At every imaging timepoint (days 0–63), we scored six binary wound-response outcomes on each *Porites* fragment from bright-field and fluorescence images: *coenosarc coverage* (complete coenosarc covering the original wound area, yes/no); *regenerated* (fully formed polyps reappearing in the center of the wound bed, yes/no); *algal plug* (dense algal plug occupying the wound bed, yes/no); *yellow aggregations* at the wound margin or in wound (yes/no); *pink aggregations* at the wound margin (yes/no); and *red fluorescent protein (RFP) signal* at the wound margin (yes/no).

#### Analysis

We modelled each of the six outcomes separately with a generalized linear mixed model (binomial error, logit link) using glmmTMB (Brooks et al. 2017) in R version 4.5.2 (R Core Team 2025). Fixed effects were treatment (airbrush, scrape, airbrush + scrape), time, and the treatment × time interaction. Time was modelled as a natural cubic spline with three degrees of freedom (to capture non-linear temporal patterns). We included colony identity (parent genotype, n = 15; each colony contributed a single wounded fragment that was imaged at multiple timepoints) as a random intercept to accommodate within-colony correlation across time. Time was mean-centered to improve convergence. Latent-scale intra-class correlations of the colony random intercept varied substantially across outcomes (coenosarc coverage: ICC = 0.59; polyps in center: 0.38; algal plug: 0.29; pink: 0.37; yellow aggregations: 0.15; RFP: 0.003), indicating that between-colony variance is the dominant source of unexplained variation for healing outcomes but contributes little for the wound-edge fluorescence signal.

Five outcomes (complete coenosarc coverage, complete regeneration, algal plug, yellow aggregations, and RFP) exhibited complete separation — one treatment reached 100% response while another remained near 0% — which renders the maximum-likelihood estimates undefined. For these outcomes we fitted a penalized binomial GLMM with a weakly informative Normal prior on the fixed effects (Normal(0, 3) on slope coefficients, Normal(0, 10) on the intercept; Gelman et al. 2008) on the real 0/1 response. The prior shrinks coefficients gently toward zero, yielding finite identifiable estimates while leaving the binomial likelihood and the data unmodified. The pink-aggregations outcome was free of separation and used a standard binomial GLMM.

Because each outcome tested an *a priori* directional hypothesis, unadjusted p-values from Wald z-tests on the spline coefficients are the primary inference; we report Benjamini–Hochberg false-discovery-rate (BH-FDR) and Bonferroni-corrected p-values across the six tests as defensive sensitivity checks. The six outcomes (coenosarc coverage, polyps in center of wound, algal plug, yellow aggregations, pink aggregations, and red fluorescence) were specified as the cellular indicators of healing and regeneration, fixing the family of tests in advance of analysis. Model adequacy was assessed with DHARMa (Hartig 2024). Residual diagnostics for all six models are shown in Figure S4.

### Experiment 3: Microscopic evaluation of scraped and airbrushed wounds

Four fragments (3-5 cm diameter) were collected from each of ten large *Porites* colonies along the Northwest corner of Mo’orea during the austral winter of 2025. Fragments were transported back to Gump station where their bases were leveled with a bandsaw (Gryphon C-40 Aquasaw) and glued to labeled tiles. Corals were acclimated, wounded, and housed as in Experiment 2. Corals were wounded a single time using scraping or airbrushing, with wound size standardized to 8 mm diameter circles. Ten corals (one from each colony) were processed for histological examination at D2, 7, 14, and 21 by squaring off a ∼1 cm piece around the wound with a bandsaw. Sawed fragments were gently cleaned with a plastic pipette to minimize disturbance to the algal plug if present. Cleaned fragments were immediately placed in 10% neutral buffered formalin (NBF) for 16 hours, then rinsed three times with phosphate buffered saline (PBS), then three times with 70% ethanol for 15 min each and finally stored in 70% ethanol until decalcification. Sections were decalcified with CalEx-II until bubbling ceased (∼24 hours) and rinsed 3 times in deionized water and 70% ethanol. Decalcified sections were processed in isopropanol and embedded in paraffin blocks. 5 µm sections were stained with Hematoxylin and Eosin (H&E) or Masson’s Trichrome for cellular analysis.

## RESULTS

### Experiment 1: Extent of tissue regeneration in three stony coral taxa varies by genera and wound type

Across all three coral taxa, scrape injuries (which removed tissue and damaged skeleton) regenerated more completely than those created using an airbrush (which removed tissue but did not damage the skeleton). However, the extent of regeneration varied across genera: only a single *Pocillopora* fragment fully regenerated tissue-scrape wounds during the four-week observation period, while all *Acropora* and *Porites* samples exhibited complete regeneration (Figure 2A-B). Following airbrush wounds, only one coral of 11 fully regenerated after 28 days (Figure 2B) – the one regenerated coral was a *Porites*. Differences in the extent of regeneration were strongly correlated with macroalgal colonization of the wound bed (Figure 2A-B). For example, almost every airbrushed wound made in *Acropora* and *Porites* was colonized by algae in one week, and the algal plug persisted until the end of the experiment. In *Pocillopora*, all airbrushed wounds were also colonized by algae in a week, but algae only persisted in 40% of the fragments after four weeks (Figure 2B). Interestingly, while scrape wounds were resistant to algal colonization in *Acropora* and *Porites*, 60% of *Pocillopora* scrape wounds were colonized by algae (Figure 2B). Our generalized linear mixed-effects model, which accounted for paired fragments from the same parent colony and repeated measures through time, supported these patterns statistically. Airbrush wounds were colonized by algae and rarely regenerated; scrape wounds did the opposite. Of the 11 fragments in each wound type, nine scrape versus one airbrush had fully regenerated (polyps in wound center), eight scrape versus one airbrush had fully healed (complete coenosarc closure) by D28, and algal cover developed on all 11 airbrush fragments (8 of 11 still had algae at D28) versus four of eleven scrape wounds (1 of 11 by D28). Coenosarc healing followed the same wound-type pattern: scrape wounds had roughly 26-fold higher odds of complete coenosarc closure than airbrush wounds (binomial GLMM, OR = 26.0, 95% CI 3.3–202, P = 0.002; Firth penalized refit OR = 17.3, 95% CI 3.2–93, P = 8.9 × 10⁻⁴). Together, these results show that species and skeletal architecture of the wound, rather than parent colony identity, were the dominant factors shaping whether colonies achieved full tissue regeneration.

**Figure 2.**
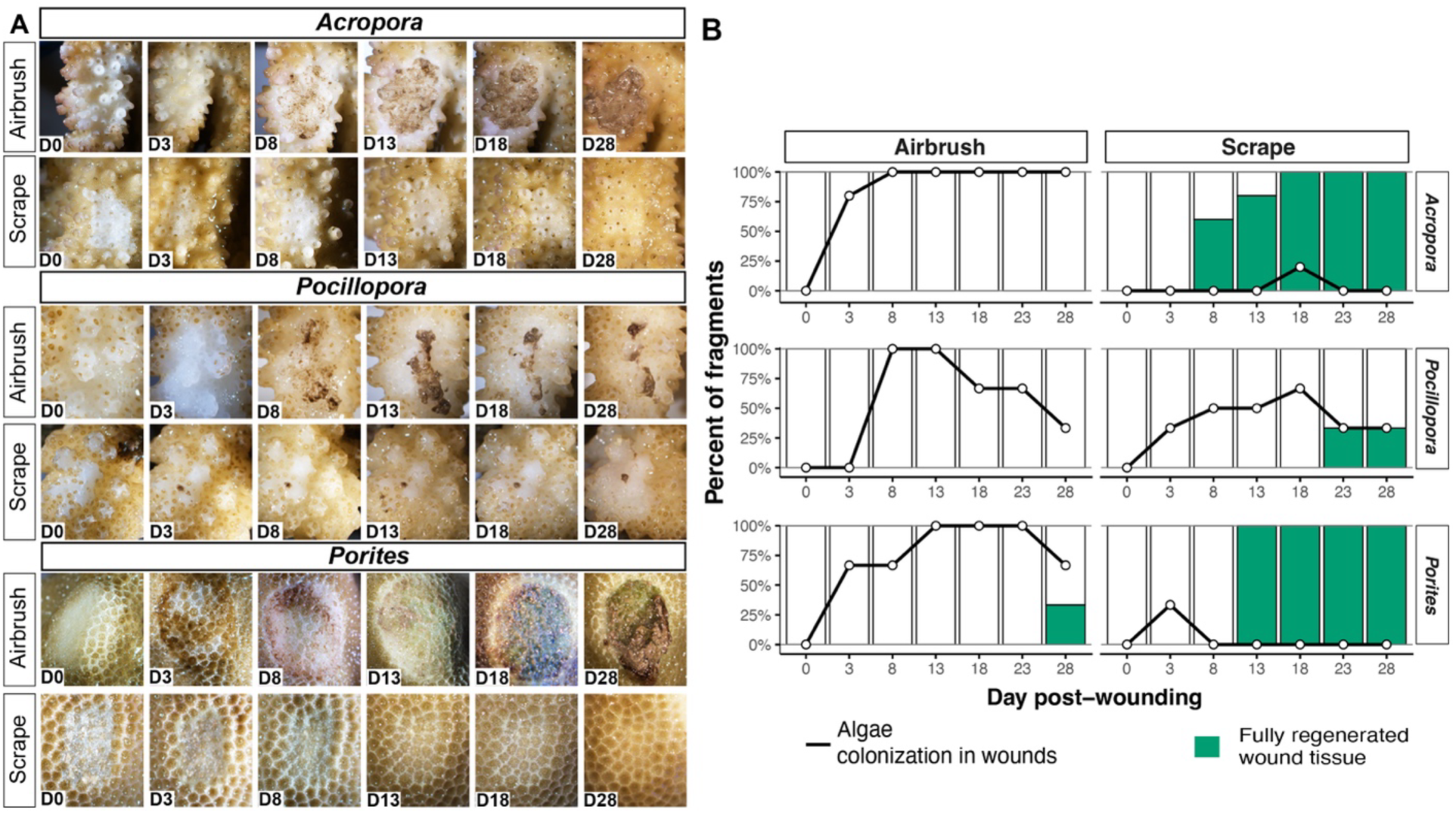
Extent of tissue regeneration in three stony coral taxa varies by genera and wound type. **(A)** Representative images of tissue regeneration in three coral genera (*Porites*, *Acropora*, and *Pocillopora*) over a 28-day period, for two injury types: airbrush (deep tissue removal, no skeletal damage) and scrape (tissue and skeletal damage) (see Methods for Experiment 1). Columns represent time post-wounding (D0, 3, 13, 18, and 28). D0 indicates the initial state immediately after standardized tissue removal (wound creation), while subsequent days track changes in wound morphology, algal colonization and tissue regeneration. Colonies were photographed *in situ* under ambient light conditions to document wound closure and tissue re-growth. **(B)** Green bars show the proportion of fragments that completely regenerated tissue across the wound bed through time for *Acropora*, *Pocillopora*, and *Porites* under two wound types (scrape and airbrush). Black solid lines represent the proportion of fragments whose wounds were colonized by algae.

Algal colonization showed an inverse pattern to tissue regeneration (Figure 2B). Airbrush wounds were colonized by algae far more often than scrape wounds: the model-predicted probability of colonization was 68–79% across genera under airbrushing but only about 5–8% under scraping, giving scrape wounds roughly 16-fold lower odds of algal colonization (Firth penalized logistic regression, OR = 0.062, 95% CI 0.027–0.144, P < 0.001). Wound type dominated this outcome (drop-one χ² = 20.94, df = 1, P < 0.001) with no detectable species effect (χ² = 0.32, df = 2, P = 0.85); the wound type × species interaction was statistically significant (χ² = 9.28, df = 2, P = 0.0096) but reflected differences in magnitude, not direction: per-genus Firth ORs (Scrape vs Airbrush) preserve the protective effect of scraping across all three genera, although the magnitude varies (*Acropora* OR = 0.010, 95% CI 0.001– 0.062; *Porites* OR = 0.036, 95% CI 0.005–0.256; *Pocillopora* OR = 0.51, 95% CI 0.14–1.78). The directional effect is therefore consistent across genera, but its magnitude is weakest and not detectable as significant in *Pocillopora* at this sample size, which the additive model cannot resolve. Algal colonization therefore tracked wound type rather than species, and inversely mirrored regeneration.

Polyp regeneration within the wound bed followed the same pattern but more starkly. As with healing, the treatment × species interaction was not significant (likelihood-ratio test: χ² = 2.07, df = 2, P = 0.355), so we fit an additive model in which both terms strongly predicted regeneration (treatment: χ² = 31.79, df = 1, P = 1.7 × 10⁻⁸; species: χ² = 14.13, df = 2, P = 8.5 × 10⁻⁴). Scrape wounds had roughly 52- fold higher odds of full regeneration than airbrush wounds (Firth penalized logistic regression, OR = 52.2, 95% CI 9.7–281, P = 4.2 × 10⁻⁶). Relative to *Acropora*, *Pocillopora* was roughly 11-fold less likely to regenerate (OR = 0.091, 95% CI 0.021–0.405, P = 1.6 × 10⁻³), whereas *Porites* did not differ from *Acropora* (OR = 1.07, 95% CI 0.376–3.04, P = 0.90). Predicted probabilities of regeneration under scrape were 58.3% for *Acropora* and 60.1% for *Porites* but only 9.4% for *Pocillopora*, versus ≤ 1.8% under airbrush for all three genera. Regeneration was therefore effectively scrape-exclusive: across 154 scored observations only a single regeneration event occurred under airbrushing, compared with 36 under scraping.

### Experiment 2: Deep tissue loss and algal colonization in Porites — macroscopic outcomes and microscopic dynamics

Based on our results from Experiment 1 showing that scraped wounds in *Porites* regenerated while airbrushed wounds were always colonized by algae, we next focused on distinguishing if the difference in healing and regeneration was due to differences in the amount of tissue removal or skeletal damage. Scraped wounds regenerated between 1-2 weeks post injury (Figure 3A-B), whereas wounds created with an airbrush (independent of the presence or absence of skeletal damage) were rapidly colonized by algae, which in some cases, remained even after eight weeks (Figure 3A-B). In all treatments, we observed yellow aggregations first at the wound margins. For the scrape-only wounds, yellow aggregations also covered the wound, whereas for the two wound types made with the airbrush, yellow aggregations were at the edge of the algal plugs but also covered newly regenerated tissue (airbrush and airbrush + scrape) (Figure 3A-D). When tissue regeneration was unimpeded by algae, these yellow aggregations persisted for at least five weeks, with small foci visible even at eight weeks (Figure 3A). Despite microscopic evaluation, the identity of these aggregations remains unresolved, although fluorescent analysis allowed us to rule out Symbiodiniaceae. In *Porites*, the contrast between scrape and airbrush wounds changed significantly through time for five of the six scored outcomes (penalized binomial GLMM, treatment × time interaction), as quantified below. Lastly, at the edge of algal plugs adjacent to newly regenerated coenosarc tissue, we observed pink-colored tissue reminiscent of pink line syndrome (PLS) (Figure 3A, pink arrows and Figure S1A-D). This pink tissue was visible using an RFP filter, and often RFP was present even when pink tissue was not visible (Figure S1A-D).

**Figure 3.**
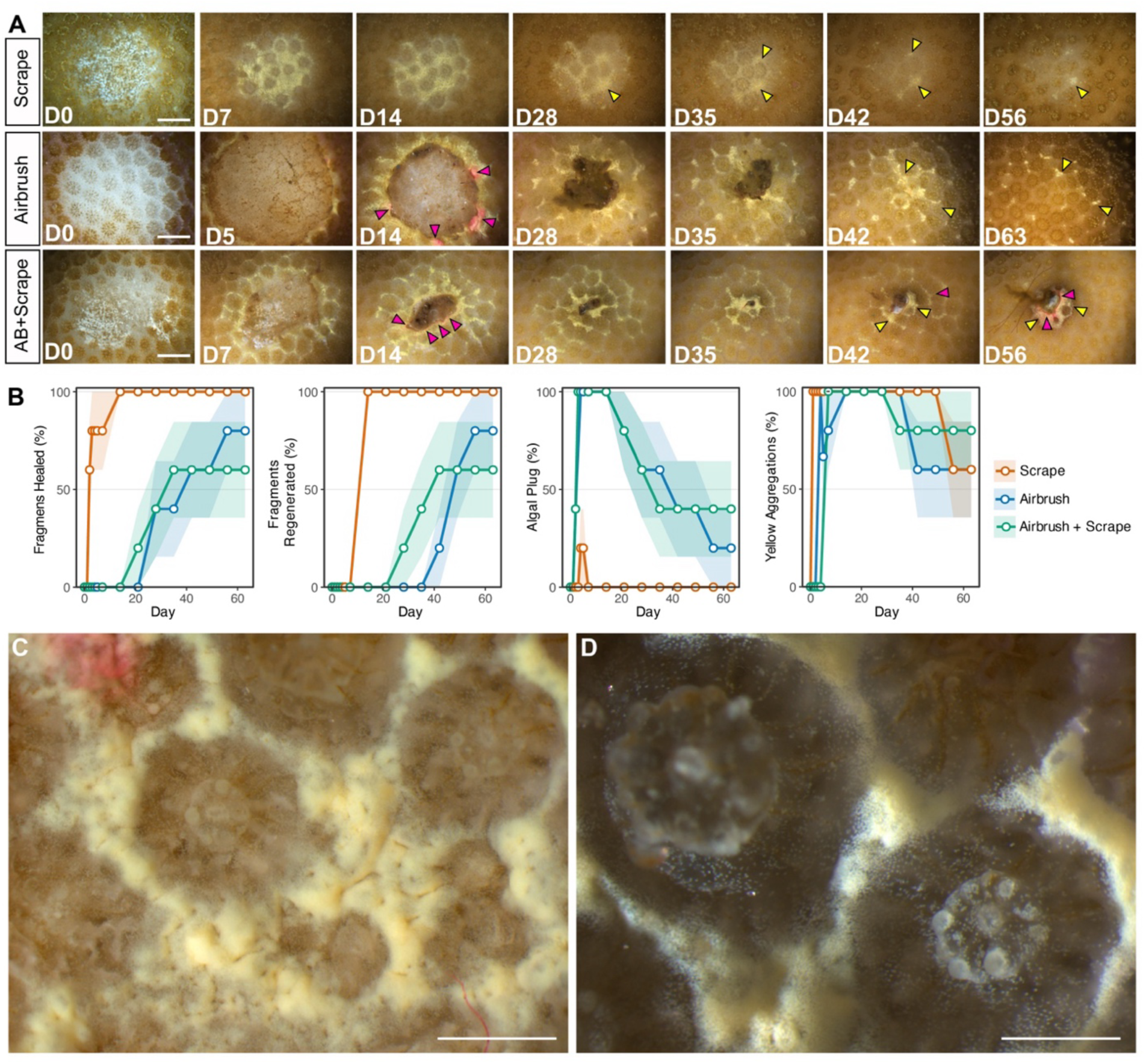
Deep tissue loss with or without skeletal damage inhibits rapid healing and allows for algal colonization in *Porites* wounds. **(A)** Macroscopic comparison of tissue regeneration following skeletal and tissue damage (scrape), deep tissue removal (airbrush) and deep tissue removal plus skeletal damage (airbrush + scrape). Algal colonization of airbrush and airbrush + scraped wounds is apparent at D5 and D7 respectively and these plugs become more distinct with time. Yellow aggregations are visible across the wound bed and examples indicated by yellow arrows. Pink tissue at the wound margin around algal plugs indicated by pink arrows. **(B)** Quantitative comparison of four key parameters during tissue healing and regeneration scored as percent of fragments: complete coverage of the wound with coenosarc, wounds that completely regenerated polyps, algal plug persistence in the wound and persistence of yellow particle aggregations around and in the wound bed. **(C-D)** High magnification images showing yellow aggregations which reveal they are individual cells. Scale bars = 1 mm.

Wound type reshaped the entire healing trajectory: treatment × time interactions were significant for five of the six tracked outcomes (Figure 3B; Table S10), with p-values ranging from 3.2 × 10⁻⁵ (RFP) to 0.021 (pink aggregations); only the presence of algae lacked a significant treatment × time interaction (p = 0.21). Scraped wounds reached 100% coenosarc coverage and 100% polyp regeneration by D14, whereas airbrush and airbrush + scrape wounds remained at ≤ 80% coenosarc coverage and ≤ 10% polyp regeneration through D63 (coenosarc coverage: p = 0.006; regenerated: p = 0.003). Some algal plugs formed transiently in scraped wounds, and all of these were gone by D14. In contrast, algae appeared in 100% of the two airbrush treatments, and 20-40% of these corals still had algae in their wound beds at D63. This stark contrast in the temporal dynamics among treatments conflicts with the lack of a significant treatment × time interaction (p = 0.21), likely due to the non-monotonic trajectories which weren’t captured by the spline model despite the clear effect of wound type on algal dynamics. Higher-order splines did not rescue the interaction: refitting with 4 and 5 spline degrees of freedom gave joint Wald p-values of 0.15 and 0.27 respectively (Table S12), indicating that the treatment × time interaction is not detectable at the per-treatment sample sizes available regardless of spline flexibility. The visual contrast between treatments is therefore driven by main effects and marginal trajectories rather than a formal interaction.

The significant treatment × time interaction for RFP (p = 3.2 × 10⁻⁵) arose because the RFP signal attenuated in scraped wounds by ∼ D30 coincident with coenosarc closure, but persisted in 50–100% of the fragments along the wound margin in the airbrush and airbrush + scrape treatments for the full observation period. The presence of RFP is consistent with sustained injury signaling at the algal–tissue interface. Yellow aggregations appeared early across all treatments and persisted in roughly half to all fragments through D63 (p = 0.012). Pink aggregations were variable but rose to ∼ 50% prevalence in scraped wounds late in the trajectory while remaining lower and more variable in airbrush treatments (p = 0.021). Together, these results show that deep tissue removal, with or without skeletal damage, reshapes the *Porites* healing trajectory by enabling persistent algal occupation of the wound bed, leading to ongoing healing at the tissue margin and a near-complete failure of polyp regeneration over the 63-day observation period.

### Tissue remnants at the injury site provide for rapid coenosarc coverage of the wound bed

To better understand and quantify early changes in cellular coverage of the wound bed, we investigated the first five days post injury (Figure S2) in scrape wounds where the wound bed was not colonized by algae (Figure 4A-H’). Owing to fluorescence of coral tissue in *Porites*, we also used two, dual bandpass filters to observe native GFP and RFP signals during tissue healing (Figure 4A’-E’, F-H’). Observing corals immediately after injury showed damaged skeleton with open spaces between the translucent skeleton in the wound center (Figure 4A-A”). After 24 hours, tissue was visibly beginning to populate the spaces between the skeleton (Figure 4B-B”). Between 48 and 72 h post wounding, the wound bed was completely covered with a thin layer of new tissue and yellow aggregations (Figure 4C-C”). Following coverage with new tissue, GFP fluorescence revealed newly forming skeleton across the wound bed coincident with the original pattern of damaged calyxes surrounding polyps (Figure 4D-E” and yellow arrows). An RFP signal appeared at the margin of the original wound ∼72 h post injury (Figure 4F, F’) and rapidly disappeared 24 h later (Figure 4F-G). In addition to the transient RFP signal at the original wound margins, we did observe persistent RFP expression around places where algae occasionally colonized wound beds and in areas where we noted pink tissue (Figure 4F–H and Figure S1).

**Figure 4.**
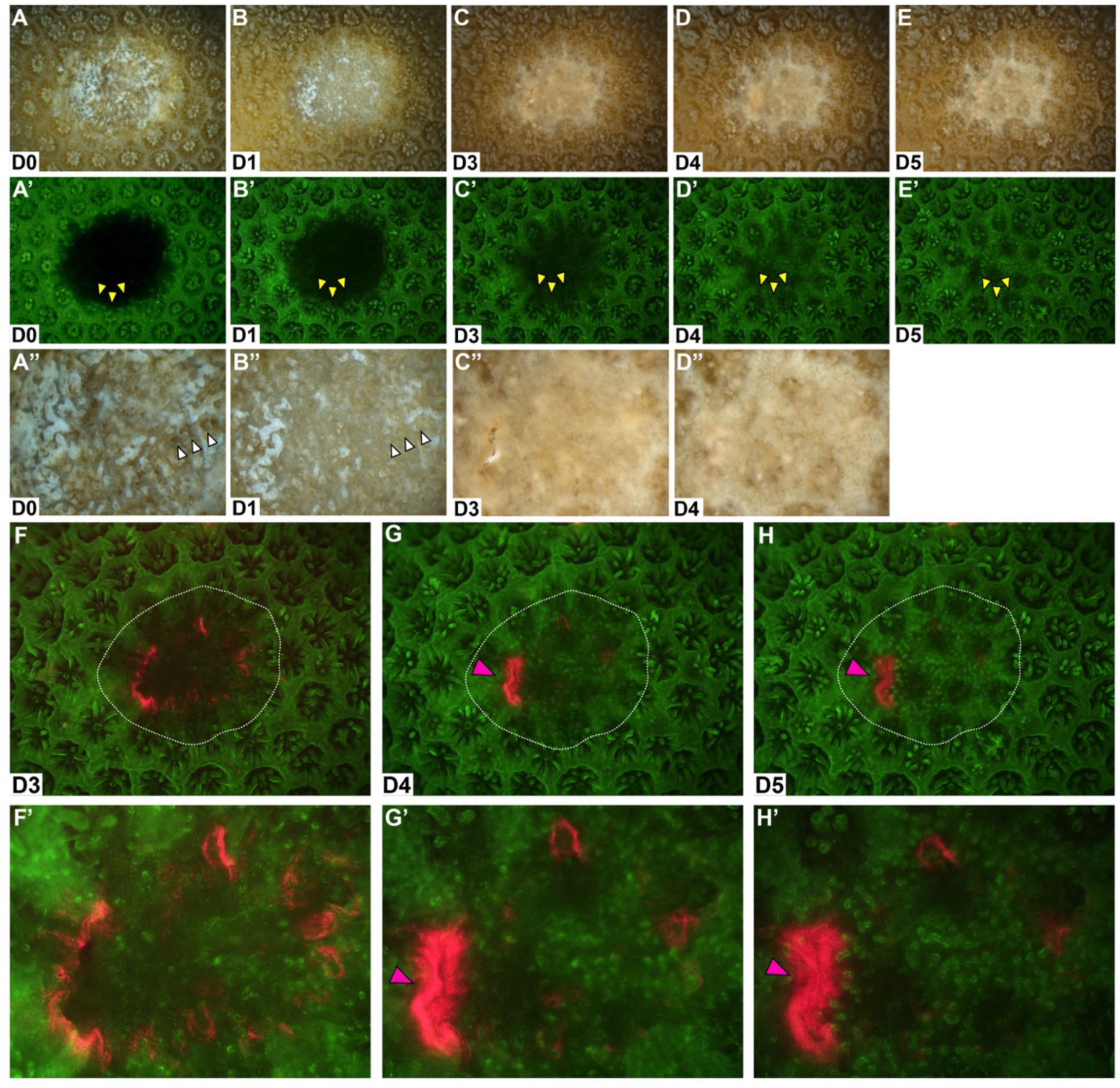
The wound healing phase of tissue regeneration is completed between 48-72 h post injury in scrape injuries. (A-E”) Whole mount photomicrographs of *Porites* wound beds in brightfield (A-E), native GFP (A’-E’) and high magnification of the wound center depicted in A-E (A”-E”). Immediately following a scrape injury, the wound center contains damaged skeleton and tissue with porous spaces visible between the translucent skeleton (A, A”, white arrows). 24 h later, cells are visible emerging from within the previous empty spaces (B, B”, white arrows) and by 72 h post injury the wound bed has a thick covering of cells and yellow aggregated spots (C, C”). Between D4-5 post injury, these cells begin to segregate around newly forming calyx cups (D, D”, E, E”). Native GFP in the wound bed reveals new cells atop the newly forming skeleton (A–E’ and yellow arrows). Yellow arrows depict reformation of three calyx skeletal cups around existing polyps. **(F-F’)** Native RFP imaging reveals a transient signal around the edge of leading-edge coenosarc where the entire multi-tissue structure is reforming (see Figure 1E for cell layers). **(G-H’)** In several areas we see persistent RFP similar to that observed around algal plugs (see Figure S1).

### New polyps regenerate approximately one week after complete wound healing

To specifically track the temporal pattern of polyp regeneration across wound types, we used native GFP autofluorescence (Figure 5). Seven days after injury, the central wound beds of scraped wounds were covered with new tissue but lacked new polyps (Figure 5). Seven days later at D14, we observed complete polyp regeneration across the wound bed (Figure 5, white arrows indicate regenerated polyps). Although algal plugs colonized the center of airbrush and airbrush + scrape wounds, we still noted tissue healing around the margins of the mat which could be visualized by RFP autofluorescence (Figure 5). Interestingly, in the area between the original wound margin and wound edge surrounding the algal plug, we noted new polyp formation after approximately seven days mimicking the timeline for scrape wounds (Figure 5, white arrows). Consistent with this timeline, as tissue healing progressed and the algal plug area decreased within the airbrushed wound beds, we noticed that outside the encroaching area, rows of new polyps emerged, again, seven days later (Figure 5, yellow arrows). These data in *Porites* demonstrate that polyp regeneration occurs approximately one week after coenosarc has completely healed tissue wounds (Table S11).

**Figure 5.**
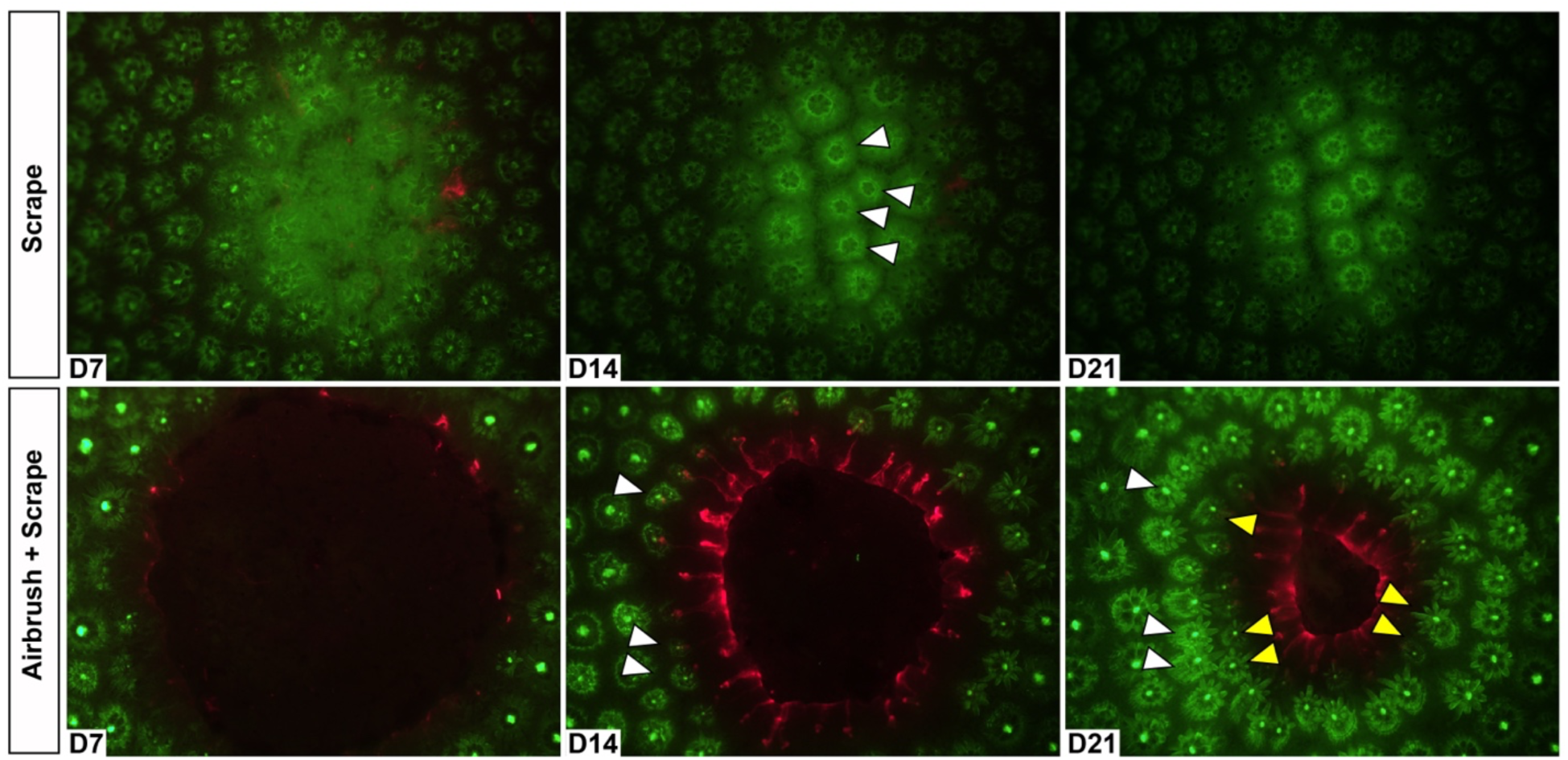
Polyp regeneration occurs approximately nine days after complete coenosarc regeneration. In scraped wounds where algal colonization does not occur, newly regenerated polyps are visible with the center of the wound bed (white arrows in top row). RFP fluorescence is visible at the leading edge of regenerated coenosarc surrounding the algal plug in airbrush + scrape wounds, and polyps are regenerated just behind this leading edge (white arrows in bottom row). This pattern persists as the area of algal colonization decreases and the healing front of new coenosarc progresses towards the center. Yellow arrows indicate a new row of regenerated polyps which form between D14 and D21 post injury with white arrows at D21 indicating the polyps regenerated at D14.

### Experiment 3: Algal colonization of the wound bed leads to deep tissue destruction and skeletal degradation

In histological sections from scrape wounds, we observed cells and yellow aggregations at D2 (Fig. 6A). By D7 post injury, yellow aggregations and new polyps were visible atop the wound bed (Fig. 6B green and yellow arrows respectively). Moreover, at D7 and D14, the deep tissue architecture outside and inside the wound margins was indistinguishable (Fig. 6B-C). Airbrush wounds exhibited a markedly different progression (Fig. 6D-G). Two days post injury (D2) cellular remnants were scattered throughout the wound bed beneath the algae (Fig. 6D). For the most part, algal plugs did not survive histological processing, but an empty depression was visible in the wound bed where the algal plug had been located. Figure 6G provides one example where algae are still visible atop the wound bed. Sections prepared at D7, 14 and 21 post injury, show that cellular remnants were no longer visible in the deeper skeleton. Instead, the deep endolithic algae layer progressively migrated up towards the wound-bed (Fig. 6E-G). Yellow aggregations were dense at the wound margins (6F-G). Surprisingly, we observed that the skeleton progressively disintegrated over time beneath the wound bed and coral cells did not appear to migrate into the wound bed from the wound margins (Fig. 6E-G). Together, our observations suggest that algal colonization of airbrush wounds inhibits healing and regeneration. Importantly, these data also suggest that algae leads to progressive loss of tissue around the deeper skeleton, which is followed by degradation of the deeper skeleton directly beneath the wound area.

**Figure 6.**
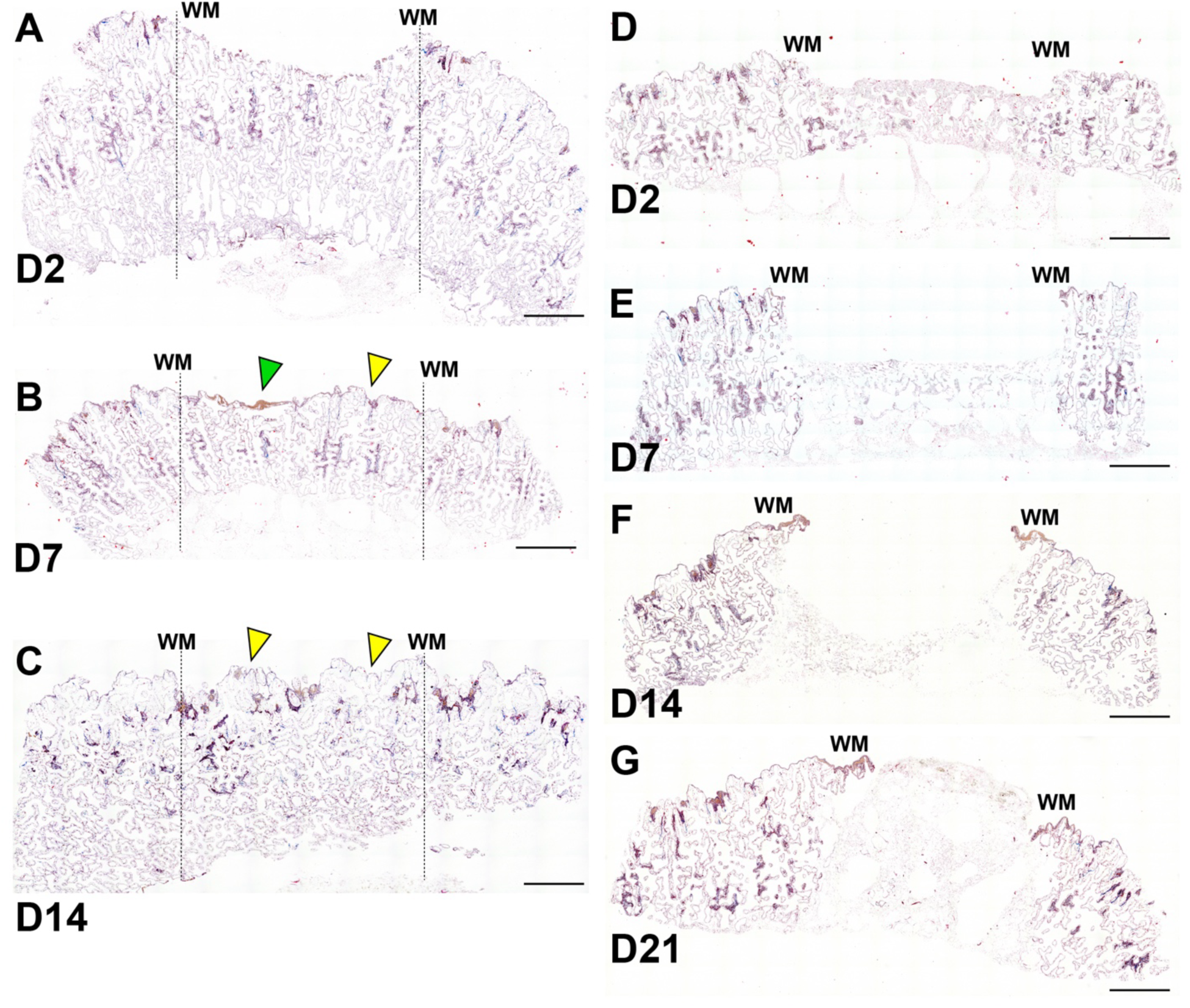
Histology. Algal colonization of the wound bed leads to deep tissue destruction and skeletal degradation. Tissue sections stained with Masson’s Trichrome prepared from healing *Porites* fragments wounded by scraping (A-C) or airbrushing (D-G). **(A)** Two days post injury (D2) in scrape wounds, new tissue is visible atop the coral skeleton in the wound bed and intact tissue is visible throughout the deeper skeleton beneath the wound bed. **(B-C)** At D7 and D14 post injury yellow aggregations and new polyps are visible in the wound center (green and yellow arrow respectively). **(D)** 48 h post injury (D2), airbrushing has removed most tissue atop and within the deeper skeleton in the wound bed. **(E-G)** From D7-D21, the presence of algae in the wound bed prevents tissue at the margins from moving across the wound and the underlying skeleton shows signs of degradation. Although the algal plugs did not normally survive tissue preparation, algae is visible atop the wound bed in (G). WM = wound margins. Scale bars = 200 µm.

## DISCUSSION

### Depth of tissue loss, not skeletal damage, gates regeneration

Regenerative capacity in *Hydra*, *Nematostella*, and *Hydractinia* has mostly been studied under ideal laboratory conditions where polyps are isolated from competing organisms. While the collective body of work using these models has produced a deep understanding of regeneration at the cell and molecular level, far less is known about regenerative ability in these single, free-living polyps under more natural conditions. In contrast, regenerative ability in colonial cnidarians whose polyps are networked within a calcium carbonate skeleton (stony corals) lack equivalent cellular-to-organismal characterization in natural or laboratory conditions. Stony corals in a reef environment must repair tissue damage in a community context where macroalgal competitors are ever-present.

In this study, we found that regenerative ability was present in species from three coral genera, but this capacity was gated by the wound environment, specifically by whether algal competitors occupied the exposed skeleton before tissue healing could occur. Importantly, we observed that depth of tissue removal — not skeletal damage — sets this contingency. Recognizing this distinction frames regeneration in colonial, calcifying cnidarians as a problem where ecology and molecular biology collide.

How deeply tissue was removed, and whether algae then colonized the wound bed, governed coral healing and regeneration far more than did skeletal damage, and its consequences depended on tissue and skeletal architecture of the wounded species. During our four-week observation period, we saw tissue healing and regeneration after scrape injuries in all *Porites* and *Acropora*, but only rarely in *Pocillopora*, whereas airbrush wounds were colonized by algae, which appeared to inhibit healing and regeneration. Key differences in skeletal architecture and tissue organization further influenced healing and regeneration. *Porites* spp. have closely packed polyps connected to deep tissue reserves (∼5 mm) within a porous skeleton (Barnes and Lough 1992; Uribe et al. 2025) and commonly house a community of endolithic algae within their skeleton just below the living coral tissue (del Campo et al. 2017). In contrast, *Acropora* and *Pocillopora* have a thin veneer of tissue over a dense skeleton, with more sparsely dispersed polyps separated by a greater amount of coenosarc. *Acropora* have deep-set corallites, which house each polyp, whereas the polyps in *Pocillopora* sit in shallow corallites (Veron and Pichon 1976; Glynn 1999).

The architectural differences between *Porites* (perforate), and *Acropora* and *Pocillopora* (imperforate) and between *Acropora* and *Pocillopora* (deep vs. shallow corallites) appear to influence their respective regenerative responses. *Acropora hyacinthus* achieved rapid, uniform healing after scrape wounds, with all wounds reaching closure between D13-28. Over half of the scraped corals showed pigmented polyps in the wound bed by D3, suggesting that these polyps may have been able to evade significant damage from the scrape treatment, thus potentially serving as tissue reservoirs for coenosarc healing and polyp regeneration that led to rapid lateral growth. This supports the important contribution of residual tissue to facilitate rapid healing. By contrast, airbrushing treatments damaged polyps more completely, which removed the tissue reserves provided by the deep corallites and led to a delay in wound recovery and regeneration. Rapid healing from spared tissue in *Acropora* aligns with its opportunistic life history that prioritizes rapid growth (Bak 1983; Pisapia et al. 2016) where this strategy may be effective for frequent branch loss or minor injuries but may be less resilient to complete polyp loss from certain types of predators.

Deep tissue reserves protected *Porites* against scrape wounds but left it vulnerable to airbrush injuries. With deep corallites, the genus demonstrated robust healing from scrape wounds, achieving coenosarc healing and polyp regeneration within two weeks or less. As was seen in our *Acropora* corals, deep corallites likely protected polyps from scrape injuries. *Porites* tissue and skeletal architecture conferred additional resilience by having a perforate skeleton, in which tissue persisted deep into the skeleton and could help facilitate rapid healing in the wound bed. A porous skeleton, however, allowed airbrush wounds to damage these deep tissue reserves, and created an extensive network of open space within the skeleton for algal colonization. The extended lag in airbrush wound regeneration, which remained unhealed through D28, underscores how algal competition can prevent healing and regeneration, although the extent to which this inhibition is physical, chemical or both remains unresolved.

While several non-mutually exclusive mechanisms might explain how algae in the wound bed impedes tissue healing, deciphering the relative contribution of turf-algae and endolithic algae is required to resolve the precise dynamics of tissue regeneration under algal influences. For example, microbial activity associated with turf-algae can consume O_2_ and create a hypoxic environment in the diffusive boundary layer at the algae-coral interface (Jorissen et al. 2016). Evidence suggests that local hypoxia at coral–algal contact zones disrupts tissue healing analogous to an infected wound (Barott et al. 2009, 2012). Algal exudates can shift water-column chemistry toward copiotrophic and potentially pathogenic bacteria (Haas et al. 2016; Wegley Kelly et al. 2022). At the local scale, contact with turf algae restructures the coral microbiome (Pratte et al. 2018), and algae specifically shift *Porites* microbiomes in Mo’orea (Brown et al. 2019). This effect can operate at multiple spatial scales (local contact plus site-level macroalgal cover) and therefore implies diffusible chemical or microbial mediators rather than purely physical contact might impede tissue healing (Briggs et al. 2021). Moreover, endolithic algal colonization of wounds in our study adds to the complexity of these physical and chemical influences. Cellular stress increases coral respiration (Paradis et al. 2019), whereas endolithic algae like *Ostreobium spp.* produce O_2_ in a hyperlocal environment that can be rapidly consumed by respiring coral tissue during daytime conditions (Kühl et al. 2008). *Ostreobium spp*. may also transfer photoassimilates to coral tissue, which might support coral healing under stress conditions (Fine and Loya 2002; Sangsawang et al. 2017). Simultaneously, rapid endolithic algae growth can weaken the skeleton through bioerosion that outpaces tissue growth (Uribe et al., 2025). Although our experiment cannot distinguish between these mechanisms, future studies pairing wound-bed O_2_ microsensors and irradiance probes, microbiome sequencing, and exometabolomics could be informative to resolve algal-coral interface dynamics under wounded conditions.

Shallow corallites and detachable polyps make *Pocillopora* the most vulnerable of the three species we examined. With its shallow corallites and imperforate skeleton, the genus was the slowest of the three genera to completely heal wounds with coenosarc, and the least likely to regenerate polyps. Unlike the other two genera, *Pocillopora* also have polyps that can detach themselves from the skeleton and leave the colony under stress (Fordyce et al. 2017; Chuang et al. 2021; Schweinsberg et al. 2021), further limiting the amount of surviving tissue available to assist with healing and regeneration. We also found that *Pocillopora* was more susceptible to algal colonization. After four weeks, 50% of scrape wounds possessed algae compared to none in either *Acropora* or *Porites* suggesting a reduced ability to reestablish an epithelial barrier in the wound bed. These factors may explain why *Pocillopora* heal and regenerate more slowly. Independent field experiments corroborate this differential algal susceptibility: *Pocillopora* suffers more severe and persistent damage from allelopathic macroalgae than *Porites* in Fijian field trials, with damage continuing to expand after algal contact was removed (Bonaldo and Hay 2014), indicating that the algal insult is an active chemical effect rather than passive co-occurrence. Together, these results support that diversity in skeletal architecture across species can affect tissue healing and regenerative ability, where species with perforate skeletons or deeply retracting polyps have access to tissue reserves that can rapidly initiate wound healing and in turn facilitate tissue regeneration.

### Algal colonization of wounds inhibit tissue healing through multiple non-exclusive mechanisms and leads to deep skeletal degradation

Our detailed microscopic evaluation of tissue healing and regeneration in *Porites* allowed us to more explicitly test several alternative hypotheses posited to explain variation in tissue repair. Our cellular analyses provide support for two complementary mechanisms. Scraped wounds healed rapidly and completely compared to airbrushed wounds due to tissue that survived below the wound. Our analysis supports that rapid coenosarc healing limits algal colonization of the wound bed. Red fluorescence appeared in scrape wounds after epithelial cells covered the wound bed and appeared to coincide with the leading edge of regenerating gastrodermis as it proceeded towards the wound center. After the coenosarc regenerated across the wound bed, red fluorescence disappeared setting the stage for regeneration of polyps approximately one week later. In contrast, airbrush wounds removed deep tissue in the skeleton beneath the wound providing time for algae to colonize and establish in the wound bed.

Our results also provide support for the hypothesis that algae impede coral healing and regeneration. Once an algal plug formed in the wound bed, healing and regeneration appeared to be delayed until the algae was replaced with coral tissue. The persistent RFP signal in coral tissue at the margin of the algal plug suggests this signal coincides with active healing or a stress response that arises where coral tissue encounters algae. Our histological data supports this conclusion as we did not observe tissue beneath the algal plug. Moreover, as the leading edge of the coenosarc moved forward coincident with reduction of the algal plug, we observed regeneration of new polyps behind this front in a similar timeline (one week) to that observed in scrape wounds. While our experimental results support that residual tissue beneath the wound bed facilitates rapid healing and reduces algae colonization, our data does not support the hypothesis that skeletal damage facilitates regeneration, as previously suggested (Lock et al. 2022). We found that algal plug formation increased predictably through time with no statistical difference between airbrush and airbrush + scrape treatments indicating that deep tissue removal is detrimental to wound healing independent of skeletal damage. Future studies disturbing superficial tissue in *Porites* with or without skeletal damage could more directly test this hypothesis.

Although we were unable to characterize turf communities in the wound beds, it is possible that the algal plugs which formed across species arose from different sources. *Porites* possess endolithic algae that reside beneath living coral tissue which, given greater light exposure in our airbrush wounds, might opportunistically grow towards the surface which antagonizes healing from the deeper marginal tissue (McCook et al. 2001; Titlyanov et al. 2005; Titlyanov and Titlyanova 2008). Surprisingly, we observed that the calcium carbonate skeleton beneath the algal plugs began to degrade when it was not protected by coral tissue and became increasingly colonized by endolithic algae, at least through the end of our observation period. This is consistent with the bioeroding nature of *Ostreobium* as it colonizes a new area of skeleton (Tribollet 2008; Tandon et al. 2023). Together, our results suggest that endolithic algae and turf algae prevent coenosarc healing and new polyp regeneration, although when colonizing algae recedes, healing and regeneration proceed in the stereotypical timeline we describe in the absence of algae supporting that this response is driven by tissue at the surface rather than beneath the wound.

### Red fluorescence coincides with coral tissue healing and stress

Our results also provide insight into the dynamics of pink lesions, red fluorescence, and melanin in response to algal colonization in *Porites*. Pink lesions exhibit peroxide scavenging abilities, and their appearance has been used as a proxy for oxidative stress or immune pathway activation (Palmer et al. 2009; Meng et al. 2025). Although red fluorescent proteins can co-exist with chromoproteins in pink lesions, the two protein groups may have distinct roles in coral immunity (Suzuki et al. 2024). While the appearance of pink lesions was variable across samples, they were often associated with wound edges surrounding algal plugs. Although RFP was almost always coincident with pink tissue, we also observed a transient RFP response that was coincident with gastrodermis regeneration in scrape wounds.

Despite initial RFP signaling in all corals, the dynamics of RFP varied between wound types. Scrape wounds exhibited early, transient RFP signals followed by rapid attenuation, indicating an acute wound response and resolution. Airbrush and airbrush + scrape wounds maintained elevated RFP expression for over 60 days along the wound margin in contact with the algal plugs. Together, these dynamics are most parsimoniously explained by RFP marking active wound-edge tissue migration: the signal is transient in scrape wounds because the regenerating coenosarc leading edge rapidly converges and closes, and persistent in airbrush wounds because algal occupation prevents the leading edge from converging. This interpretation is consistent with the antioxidant and reactive-oxygen-species scavenging functions previously reported for coral fluorescent proteins (Bou-Abdallah et al. 2006; Palmer et al. 2009). We cannot, however, separate a marker-of-active-healing interpretation from a non-exclusive ROS response at the coral–algal-interface interpretation without paired molecular markers; future studies pairing RFP imaging with antioxidant or ROS assays, and with algae-free scratch controls in the same individuals, could decouple these two non-mutually-exclusive functions.

In addition to pink lesions and RFP, we also observed aggregation of yellow pigmentation throughout healing. We found that these aggregations were immediately present in all treatments and failed to disappear from over half of the samples after 60 days. In unhealed wounds, aggregations were present along the healing edge whereas in fully closed wounds, yellow pigment persisted within coenosarc but not polyps. The chronic appearance of yellow aggregations across all wound types suggests that this pigment is not associated with specific phases of regeneration, or competition with algae. In fact, our results support that the presence of these aggregations may be a useful indicator that tissue has sustained some type of damage within four weeks, although the type of damage may not be easily determined. Coral regeneration studies would benefit from clarity about the source and function of these yellow (pigmented) cells which are obvious on living reefs and suggest recent damage independent of the source.

## Conclusion

Our study shows that coral wound healing and regeneration are affected by complex interactions between resident tissue and biotic competitors, architectural features of the skeleton, and genus-specific factors that require further investigation. We found that rapid epithelial repair can occur in coral species with perforate skeletons and deep tissue reserves, but also in some species with imperforate skeletons and deep corallites, while another species with an imperforate skeleton and shallow corallites is slow to heal. These morphological correlates of healing and regeneration suggest that coral resilience to chronic disturbance depends, in part, on physical features of the affected coral species but also on species-specific factors. Our microscopic characterization of wound healing in *Porites* revealed the biphasic nature of tissue regeneration in stony corals which is consistent with recent general characterization of regeneration across metazoans comprising a healing phase followed by transition to a phase of tissue morphogenesis (growth and patterning of functional replacement tissue) (Seifert et al. 2023). We observed that rapid barrier restoration between wound edges is required to proceed to coenosarc healing. Once coenosarc healing is complete, polyp regeneration can proceed in a remarkably stereotypical progression at the margins of receding algal plugs. However, slow progression of either healing stage allowed for opportunistic algae to colonize the wound bed which in our study was clearly antagonistic to further subsequent regeneration. Yet, as algal plugs receded in some wounds, local inhibition of tissue healing, be it physical or chemical, was relieved. This speaks to the high, endogenous regenerative ability of corals and supports the need to further investigate how this information might be used to coax regeneration in damaged reefs. Our study re-frames cnidarian regeneration in colonial, calcifying species as an ecological problem layered on top of a cellular one: the regenerative capacity is present, but whether it expresses depends on a race between coral tissue and competing algae on the wound bed.

The ecological relevance of our findings requires validation under field conditions where wounds vary in size, shape, and environmental context. Temperature stress, sedimentation, and water flow can modulate healing rates and may alter the relative response of different species to different wound types. Understanding regenerative mechanisms, and the interactions among those mechanisms, wound types, and environmental factors will be essential for predicting coral resilience and persistence under global change. Regeneration in reef corals is therefore not a fixed species property but a contingent outcome, set by how deeply an injury removes tissue and whether algae can colonize exposed skeleton. This distinction matters for reef futures, because corallivores and disturbances differ systematically in the depth of tissue they remove — surface scraping by parrotfishes versus the deeper tissue loss inflicted by gastropods, seastars, and bleaching — so disturbance regimes that shift toward deep tissue removal should erode regenerative capacity disproportionately, converting transient wounds into persistent, algal-occupied lesions. Identifying which corals, and which reefs, retain the capacity to close such wounds under intensifying and interacting disturbance is now a tractable and pressing goal.

### Limitations of the current study

Although our experimental design allowed us to test regenerative ability across species, study of *Pocillopora* and *Acropora* lacked the microscopic precision deployed for Porites. Future, detailed work, will allow for general principles established in this work to be explored more deeply in these and other species. We also note that Porites in Mo’orea includes several morphologically convergent species (*P. lobata, P. lutea, P. rus*, and the *P. lichen* complex) that we could not reliably distinguish in the field and that may differ in tissue depth; the very feature on which our depth-of-removal mechanism rests. Within-genus species-level variation in tissue depth therefore remains a potential confounder for the headline mechanism, and we encourage replication of these experiments with molecular species verification (Primov et al. 2024). While we directly tested how skeletal damage affects healing by crossing our injury types, we could not directly test the hypothesis following just surface removal of tissue in Porites. While this was impossible using our current design, future experiments with *Porites* could develop a method to remove only the surface epidermis and cross that with skeletal damage to see if skeletal damage in this context does or does not affect healing and regeneration. Lastly, studying free-living animals that lack established molecular resources is challenging, and this limits the cell and molecular precision that can be exported from these models. Still, our tissue preparation methods do provide excellent material for deeper study and the validation of existing antibodies and reagents, or the development of new reagents, will allow for deeper investigation into the mechanisms underlying tissue regeneration and competition between algae and coral tissue.

## Acknowledgements

We thank Lily Zhao, Kai Kopecky, Madeline Pacheco, Michelle Diminuco, and Joshua Sarli for their invaluable assistance in the field and laboratory. Their careful work and dedication contributed greatly to the success of this project. We are also grateful to the staff of the Richard B. Gump South Pacific Research Station for logistical support and access to facilities, and to the people of Mo’orea for sharing their natural environment. This research was partially supported by funding from the National Science Foundation (OCE-1851510 and OCE-1851032) to A.C.S and C.W.O, from the Keck Foundation to A.C.S, C.W.O and A.W.S, from the College of Arts and Sciences at the University of Kentucky to A.W.S and from the University of California, Santa Barbara to A.C.S. All work was conducted under permits from French Polynesia (collection permit DIREN #01319, 2-4m depth).

## Data Availability

Processed data and all analysis code are available at https://github.com/stier-lab/stier-wound-type-2025-acr-por and archived at Zenodo (DOI: [PLACEHOLDER — to be minted on acceptance]). Raw images and histological sections are available from the corresponding author on reasonable request.

## Declaration of generative AI and AI-assisted technologies in the manuscript preparation process

During the preparation of this work, the A.C.S used Claude to copy edit code. A.C.S reviewed and edited the output as needed and takes full responsibility for the content of the published article.

