## Supplemental Data and Figures for "Deep tissue removal in wounds facilitates algal colonization and inhibits healing and regeneration in tropical corals"

*bioRxiv*

This supplement supports the two quantitative experiments in the main text: Experiment 1, a paired split-colony comparison of airbrush (deep tissue removal, no skeletal damage) and scrape (tissue plus skeletal damage) wounds across three coral genera (*Acropora*, *Pocillopora*, *Porites*); and Experiment 2, a longitudinal cellular characterization of *Porites* wounds. The photo-scoring scheme, data provenance, and the neighbour rule used to resolve missing observations are described in the Methods and Data Availability statement. Throughout, Scrape denotes the Dremel treatment; all statistics derive from the canonical analysis outputs.

### Contents

**Appendix A. Experiment 1 — model selection and effect sizes (Tables S1–S5)**

**Appendix B. Experiment 1 — missing-data resolution and robustness (Tables S6–S8)**

**Appendix C. Experiment 1 — model diagnostics (Table S9)**

**Appendix D. Experiment 2 (*Porites*) — statistical detail (Tables S10–S12)**

**Appendix E. Supporting figures (Figures S1–S4)**

Figure S1. Red fluorescence and pink aggregations at the coral–algal interface (Porites).
Figure S2. Early Porites wound dynamics, days 0–5.
Figure S3. DHARMa simulated-residual diagnostics, Experiment 1 GLMMs.
Figure S4. DHARMa simulated-residual diagnostics, Experiment 2 (Porites) GLMMs.

### Appendix A.  Experiment 1 — model selection and effect sizes

*Model selection, fixed-effect odds ratios, and model-estimated probabilities for the three Experiment 1 outcomes (complete healing, polyp regeneration, algal colonization).*

**Table S1.** Wound type and coral genus act additively: the effect of scraping versus airbrushing does not differ detectably among genera, so a single additive model describes each outcome. Likelihood-ratio tests compare the wound type × genus interaction model with the simpler additive model for each binary outcome. The interaction is non-significant for healing and regeneration, justifying the additive models reported in the main text; it is significant for algal colonization but reflects a difference in magnitude, not direction — scraping is protective in all three genera (per-genus odds ratios, Table S3). *Cited in Methods and Results.*

| Outcome | Term | Chi-sq | df | P |
|---|---|---|---|---|
| Healing | Treatment × Species Interaction | 3.55 | 2 | 0.17 |
| Regeneration | Treatment × Species Interaction | 2.07 | 2 | 0.35 |
| Algal colonization | Treatment × Species Interaction | 9.28 | 2 | 0.0096 |

**Table S2.** The wound-type effect is large and statistically robust, whereas the genus effect on complete healing is not. Type-II likelihood-ratio tests for wound type and genus in each additive GLMM, with Benjamini–Hochberg (BH) and Bonferroni adjustment across the six a priori main-effect tests. All three wound-type effects remain significant after the conservative Bonferroni correction; the genus effect is significant for regeneration but not for complete healing — a result that, as Table S7 shows, hinges on a single fragment. *Cited in Results.*

| Outcome | Term | Chi-sq | P (raw) | P (BH) | P (Bonf.) |
|---|---|---|---|---|---|
| healed | treatment | 16.25 | <0.001 | <0.001 | <0.001 |
| healed | species | 4.80 | 0.091 | 0.11 | 0.54 |
| regenerated | treatment | 31.79 | <0.001 | <0.001 | <0.001 |
| regenerated | species | 14.13 | <0.001 | 0.0013 | 0.0051 |
| algal colonization | treatment | 20.94 | <0.001 | <0.001 | <0.001 |
| algal colonization | species | 0.32 | 0.85 | 0.85 | 1 |

**Table S3.** Full effect sizes behind the odds ratios reported in the main text. Fixed-effect odds ratios (95% CI) for every term in the Experiment 1 additive models. Healing is reported from the standard binomial GLMM as the primary model — the 2026 re-score removed the complete separation that previously forced penalization — with a Firth penalized refit shown as a sensitivity analysis; regeneration and algal colonization are reported from Firth models because their airbrush cells contain almost no events. Scraping raises the odds of healing roughly 26-fold (binomial GLMM) and of regeneration roughly 52-fold relative to airbrushing. The final panel gives per-genus Firth odds ratios for algal colonization (scrape vs airbrush within each genus); scraping is protective in every genus — significantly so in Acropora and Porites — confirming that the significant wound type × genus interaction (Table S1) reflects a difference in magnitude, not direction. *Cited in Results.*

#### Healing — binomial GLMM (primary)

| Term | OR (95% CI) | P |
|---|---|---|
| Intercept | 0.02 (0.00–0.15) | <0.001 |
| Scrape vs Airbrush | 25.96 (3.33–202.44) | 0.0019 |
| *Pocillopora* vs *Acropora* | 0.20 (0.04–0.97) | 0.046 |
| *Porites* vs *Acropora* | 0.46 (0.13–1.66) | 0.24 |

#### Healing — Firth penalized refit (sensitivity)

| Term | OR (95% CI) | P |
|---|---|---|
| Scrape vs Airbrush | 17.30 (3.22–92.97) | <0.001 |
| *Pocillopora* vs *Acropora* | 0.24 (0.05–1.03) | 0.056 |
| *Porites* vs *Acropora* | 0.50 (0.15–1.71) | 0.27 |

#### Regeneration — Firth

| Term | OR (95% CI) | P |
|---|---|---|
| Scrape vs Airbrush | 52.21 (9.69–281.37) | <0.001 |
| *Pocillopora* vs *Acropora* | 0.09 (0.02–0.40) | 0.0016 |
| *Porites* vs *Acropora* | 1.07 (0.38–3.04) | 0.9 |

#### Algal colonization — Firth

| Term | OR (95% CI) | P |
|---|---|---|
| Scrape vs Airbrush | 0.06 (0.03–0.14) | <0.001 |
| *Pocillopora* vs *Acropora* | 1.02 (0.39–2.67) | 0.97 |
| *Porites* vs *Acropora* | 0.73 (0.27–1.94) | 0.53 |

#### Algal colonization — per-genus Firth (Scrape vs Airbrush)

| Genus | OR (95% CI) | P |
|---|---|---|
| *Acropora* | 0.010 (0.001–0.062) | <0.001 |
| *Pocillopora* | 0.51 (0.14–1.78) | 0.29 |
| *Porites* | 0.036 (0.005–0.256) | <0.001 |

**Table S4.** Predicted probabilities translate the odds ratios into interpretable per-group risks. Model-estimated probability of each outcome for every genus × wound-type cell on the response scale (95% CI). Under scraping, healing and regeneration are likely in *Acropora* and *Porites* but not *Pocillopora*, while algal colonization stays uniformly low; under airbrushing the pattern inverts — high algal colonization with near-zero healing and regeneration. *Cited in Results.*

| Outcome | Genus | Wound type | Probability |
|---|---|---|---|
| Healing | *Acropora* | Airbrush | 2.0% (0.3–13.4%) |
| Healing | *Pocillopora* | Airbrush | 0.4% (0.0–4.5%) |
| Healing | *Porites* | Airbrush | 1.0% (0.1–8.1%) |
| Healing | *Acropora* | Scrape | 35.1% (21.4–51.8%) |
| Healing | *Pocillopora* | Scrape | 9.6% (2.4–31.3%) |
| Healing | *Porites* | Scrape | 20.0% (7.7–42.8%) |
| Regeneration | *Acropora* | Airbrush | 1.7% (0.2–11.5%) |
| Regeneration | *Acropora* | Scrape | 58.3% (41.9–73.1%) |
| Regeneration | *Pocillopora* | Airbrush | 0.1% (0.0–1.5%) |
| Regeneration | *Pocillopora* | Scrape | 9.4% (2.4–30.8%) |
| Regeneration | *Porites* | Airbrush | 1.8% (0.2–12.9%) |
| Regeneration | *Porites* | Scrape | 60.1% (39.0–78.0%) |

| Outcome | Genus | Wound type | Probability |
|---|---|---|---|
| Algal colonization | *Acropora* | Airbrush | 74.9% (47.4–90.8%) |
| Algal colonization | *Acropora* | Scrape | 6.6% (1.7–23.1%) |
| Algal colonization | *Pocillopora* | Airbrush | 78.7% (48.0–93.7%) |
| Algal colonization | *Pocillopora* | Scrape | 8.1% (1.9–28.4%) |
| Algal colonization | *Porites* | Airbrush | 68.2% (32.9–90.4%) |
| Algal colonization | *Porites* | Scrape | 4.9% (0.9–22.2%) |

**Table S5.** The healing benefit of scraping is large in *Acropora* and *Porites* and weakest in *Pocillopora*. Scrape-minus-airbrush healing contrasts within each genus, expressed as differences in predicted probability rather than odds ratios because odds ratios are not estimable where airbrush healing approaches zero. *Cited in Results.*

| Genus | Contrast | Δ probability (95% CI) | P |
|---|---|---|---|
| *Acropora* | Scrape vs Airbrush | 0.33 (0.17, 0.49) | <0.001 |
| *Pocillopora* | Scrape vs Airbrush | 0.09 (-0.03, 0.21) | 0.14 |
| *Porites* | Scrape vs Airbrush | 0.19 (0.02, 0.36) | 0.027 |

### Appendix B. Experiment 1 — missing-data resolution and robustness

*An audit of the 2026 re-score, leave-one-out and missing-data sensitivity analyses, a model-free day-28 corroboration, and an ordinal confirmation of the binary healing endpoint.*

**Table S6.** An auditable record showing the missing-data re-score followed a transparent rule and that only two cells drive the change in statistical conclusions. Every observation changed in the 2026 re-score is listed with its bracketing (t−1, t+1) states, the value implied by the neighbour rule, and the value finally adopted. Most changes are straightforward neighbour-rule fills; the two cells that eliminate complete separation in *Pocillopora*-scrape healing both belong to fragment 6b — the fragment whose influence is quantified in Table S7. *Cited in Methods.*

| Frag. | Day | Outcome | t−1 | t+1 | Prior | Rule implies | Adopted | Classification |
|---|---|---|---|---|---|---|---|---|
| 6b | 28 | healed | incomplete | | | | yes | NA-fill: beyond rule (Craig=unresolved) |
| 6b | 28 | debris | no | | | | no | NA-fill: beyond rule (Craig=unresolved) |
| 1a | 23 | healed | incomplete | incomplete | | incomplete | incomplete | NA-fill: matches bracketing rule |
| 1a | 23 | debris | yes | yes | | yes | yes | NA-fill: matches bracketing rule |
| 2a | 13 | regenerated | no | no | | no | no | NA-fill: matches bracketing rule |
| 2a | 13 | healed | no | no | | no | no | NA-fill: matches bracketing rule |
| 4a | 23 | healed | incomplete | incomplete | | incomplete | incomplete | NA-fill: matches bracketing rule |

| Frag. | Day | Outcome | t−1 | t+1 | Prior | Rule implies | Adopted | Classification |
|---|---|---|---|---|---|---|---|---|
| 4a | 23 | debris | yes | yes | | yes | yes | NA-fill: matches bracketing rule |
| 6b | 13 | healed | incomplete | incomplete | | incomplete | incomplete | NA-fill: matches bracketing rule |
| 6b | 28 | regenerated | yes | | | yes | yes | NA-fill: matches bracketing rule |
| 9b | 8 | regenerated | no | no | | no | no | NA-fill: matches bracketing rule |
| 9b | 8 | debris | no | no | | no | no | NA-fill: matches bracketing rule |
| 10b | 18 | regenerated | no | yes | yes | | no | RE-SCORE (already-scored cell) |
| 6b | 23 | healed | incomplete | | incomplete | | yes | RE-SCORE (already-scored cell) |

**Table S7.** The headline wound-type effect is robust to analytical choices, but the genus difference in healing rests on a single fragment. Top: the healing wound-type odds ratio (binomial GLMM and Firth) and its significance across influential-fragment exclusions and missing-data scenarios — large and significant in every case. Bottom: day-28 Fisher exact tests within each genus, used because the day-28 GLMM is non-estimable under separation. The genus effect on healing reaches significance only when fragment 6b is excluded, identifying it as the single influential observation. *Cited in Results/Discussion.*

#### Healing wound-type effect across scenarios

| Scenario | Healing OR (GLMM) | Healing OR (Firth) | Wound-type P | Genus P | n |
|---|---|---|---|---|---|
| full (complete-case) | 26.0 | 17.3 | <0.001 | 0.091 | 151 |
| drop coral 6b | 23.5 | 15.8 | <0.001 | 0.011 | 144 |
| drop coral 7b | 27.4 | 18.1 | <0.001 | 0.23 | 144 |
| missing healed = yes (best) | 29.9 | 19.8 | <0.001 | 0.17 | 154 |
| missing healed = no (worst) | 25.2 | 16.8 | <0.001 | 0.087 | 154 |

#### Day-28 Fisher exact tests

| Genus | Airbrush healed/n | Scrape healed/n | P |
|---|---|---|---|
| *Acropora* | 0/5 | 5/5 | 0.0079 |
| *Pocillopora* | 0/3 | 1/3 | 1 |
| *Porites* | 1/3 | 2/2 | 0.4 |

**Table S8.** Collapsing the three-level healing score to a binary endpoint does not distort the conclusions. A cumulative-link mixed (ordinal) model of healing scored as none < incomplete < complete recovers the same strong wound-type effect as the binary model. The proportional-odds assumption is violated for wound type, indicating the scrape advantage is concentrated at the first transition (none → incomplete) — scraping mainly gets wounds started healing. Predicted category probabilities are given for each genus × wound type. *Cited in Methods/Results.*

#### Coefficients

| Term | OR (95% CI) | P |
|---|---|---|
| Scrape vs Airbrush | 2.96 (1.55–5.66) | <0.001 |
| *Pocillopora* vs *Acropora* | 0.48 (0.23–1.03) | 0.059 |
| *Porites* vs *Acropora* | 0.53 (0.25–1.14) | 0.11 |

#### Proportional-odds (nominal-effects) test

| Term | df | LRT | P |
|---|---|---|---|
| treatment | 1 | 11.31 | <0.001 |
| species | 2 | 2.00 | 0.37 |

#### Predicted category probabilities

| Wound type | Genus | Pr(none) | Pr(incomplete) | Pr(complete) |
|---|---|---|---|---|
| Airbrush | *Acropora* | 0.387 | 0.513 | 0.099 |
| Airbrush | *Pocillopora* | 0.567 | 0.383 | 0.051 |
| Airbrush | *Porites* | 0.544 | 0.401 | 0.055 |
| Scrape | *Acropora* | 0.176 | 0.578 | 0.246 |
| Scrape | *Pocillopora* | 0.306 | 0.558 | 0.136 |
| Scrape | *Porites* | 0.287 | 0.566 | 0.147 |

### Appendix C. Experiment 1 — model diagnostics

*Simulated-residual, temporal-autocorrelation, separation, and variance-component diagnostics for the reported Experiment 1 GLMMs. The companion residual plots are Figure S3 (Appendix E).*

**Table S9.** Diagnostic checks confirming the Experiment 1 models are well specified. DHARMa simulated residuals show no uniformity, dispersion, or outlier departures (companion plots, Figure S3); day-aggregated residuals show temporal autocorrelation, as expected for a deliberately time-collapsed endpoint model. After the re-score no model exhibits complete separation, so the Firth fits serve as bias-reduction sensitivities rather than necessities. Random-effect variances collapse to ~0 (ICC ≈ 0); the nested colony random effect is retained to honour the paired split-colony design rather than because it captures appreciable clustering. *Cited in Methods.*

#### DHARMa residual and temporal-autocorrelation tests

| Outcome | n | Uniformity P | Dispersion P | Outlier P | Temporal DW | Temporal P |
|---|---|---|---|---|---|---|
| Healing | 151 | 0.84 | 0.93 | 1 | 0.42 | 0.0038 |
| Regeneration | 154 | 0.39 | 0.91 | 1 | 0.40 | 0.0029 |
| Algal colonization | 150 | 0.58 | 0.75 | 1 | 0.75 | 0.044 |

#### Formal complete-separation test (detectseparation)

| Outcome | Complete separation | Separated terms |
|---|---|---|
| Healing | no | none |
| Regeneration | no | none |

| Outcome | Complete separation | Separated terms |
|---|---|---|
| Algal colonization | no | none |

#### Overdispersion, singularity, information criteria

| Model | Overdispersion | Singular | AIC | BIC | logLik |
|---|---|---|---|---|---|
| Additive | 1.107 | yes | 99.4 | 117.5 | -43.7 |
| Interaction | 0.815 | yes | 99.8 | 123.9 | -41.9 |

#### Latent-scale variance components and intraclass correlations

| Component | Variance | ICC |
|---|---|---|
| Parent colony | 0.00 | 0 |
| Fragment within colony | 0.00 | 0 |
| Residual (logit) | 3.29 | 1 |

### Appendix D. Experiment 2 (*Porites*) — statistical detail

*Per-outcome time-series statistics, the coenosarc-to-polyp regeneration lag, and a spline-flexibility sensitivity analysis for the Porites cellular experiment.*

**Table S10.** Wound type reshaped the *Porites* healing trajectory across nearly every scored feature. Per-outcome results from the Experiment 2 time-series models (penalized binomial GLMM, treatment × natural-cubic-spline(day) with a colony random intercept), giving the model used, the minimum treatment × time interaction p-value, and the number of significant interaction terms. Five of the six outcomes show a significant treatment × time interaction; only algal-plug presence does not. *Cited in Experiment 2 Results.*

| Outcome | Model | Result | Min. p | Sig. interactions |
|---|---|---|---|---|
| Algal Plug | Penalized GLMM (weakly-informative Normal prior; Gelman 2008; separation) | No significant interactions | 0.21 | 0 |
| Coenosarc Coverage | Penalized GLMM (weakly-informative Normal prior; Gelman 2008; separation) | 1 significant interaction(s), $p < 0.05$ | 0.0061 | 1 |
| Pink | Standard binomial GLMM | 1 significant interaction(s), $p < 0.05$ | 0.021 | 1 |
| Polyps in Center of Wound | Penalized GLMM (weakly-informative Normal prior; Gelman 2008; separation) | 1 significant interaction(s), $p < 0.05$ | 0.0033 | 1 |
| RFP | Penalized GLMM (weakly-informative Normal prior; Gelman 2008; separation) | 2 significant interaction(s), $p < 0.05$ | <0.001 | 2 |
| Yellow Aggregations | Penalized GLMM (weakly-informative Normal prior; Gelman 2008; separation) | 1 significant interaction(s), $p < 0.05$ | 0.012 | 1 |

**Table S11.** Polyp regeneration follows coenosarc closure by about a week, defining the two-phase healing-then-regeneration sequence described in the main text. For individual *Porites* fragments, the interval (days) between complete coenosarc coverage of the wound and the first appearance of regenerated polyps in the wound center, summarized overall and by wound type. *Cited in Experiment 2 Results.*

| Treatment | n | Median lag (d) | IQR (d) | Range (d) | Mean ± SD (d) |
|---|---|---|---|---|---|
| All | 12 | 9.0 | 7–12 | 0–21 | 9.2 ± 5.9 |
| Airbrush + Scrape | 3 | 7.0 | 7–7 | 7–7 | 7.0 ± 0.0 |
| Airbrush | 4 | 10.5 | 5.25–15.75 | 0–21 | 10.5 ± 9.0 |
| Scrape | 5 | 12.0 | 11–12 | 0–12 | 9.4 ± 5.3 |

**Table S12.** The Experiment 2 conclusions are not an artefact of spline flexibility. Each Porites outcome was refit with treatment × natural-cubic-spline(day, k) at k = 3, 4, and 5 degrees of freedom; the set of outcomes showing a significant treatment × time interaction is unchanged across df. *Cited in Experiment 2 Results.*

| Outcome | Spline df | Min interaction P | # interaction terms | # significant (p<0.05) |
|---|---|---|---|---|
| Coenosarc Coverage | 3 | 0.0061 | 6 | 1 |
| Coenosarc Coverage | 4 | 0.0081 | 8 | 1 |
| Coenosarc Coverage | 5 | 0.02 | 10 | 2 |
| Polyps in Center of Wound | 3 | 0.0033 | 6 | 1 |
| Polyps in Center of Wound | 4 | <0.001 | 8 | 1 |
| Polyps in Center of Wound | 5 | 0.02 | 10 | 2 |
| Algal Plug | 3 | 0.21 | 6 | 0 |
| Algal Plug | 4 | 0.15 | 8 | 0 |
| Algal Plug | 5 | 0.27 | 10 | 0 |
| Yellow Aggregations | 3 | 0.0079 | 6 | 1 |
| Yellow Aggregations | 4 | 0.0041 | 8 | 1 |
| Yellow Aggregations | 5 | 0.0024 | 10 | 1 |
| Pink | 3 | 0.014 | 6 | 2 |
| Pink | 4 | 0.046 | 8 | 1 |
| Pink | 5 | 0.12 | 10 | 0 |
| RFP | 3 | <0.001 | 6 | 2 |
| RFP | 4 | 0.0024 | 8 | 3 |
| RFP | 5 | 0.013 | 10 | 1 |

### Appendix E. Supporting figures

**Figure S1.** Red fluorescence and pink aggregations mark the coral–algal interface and the active healing front in *Porites* wounds — the basis for interpreting red fluorescent protein (RFP) as a marker of regenerating wound-edge tissue rather than a generic stress signal. Brightfield (A, B) and dual-pass RFP/GFP fluorescence (A′, B′) of scrape wounds; pink tissue at a small algal plug (A, B) is co-located with the RFP signal (A′, B′), and each fluorescence panel shares its brightfield field of view. (C) Pink aggregations and (D) red fluorescence over time, by wound type: both rise then fall in scrape wounds as the wound closes but persist along the algal-plug margin in airbrush and airbrush-scrape wounds. Cited in main-text Results/Discussion (Figure 4).

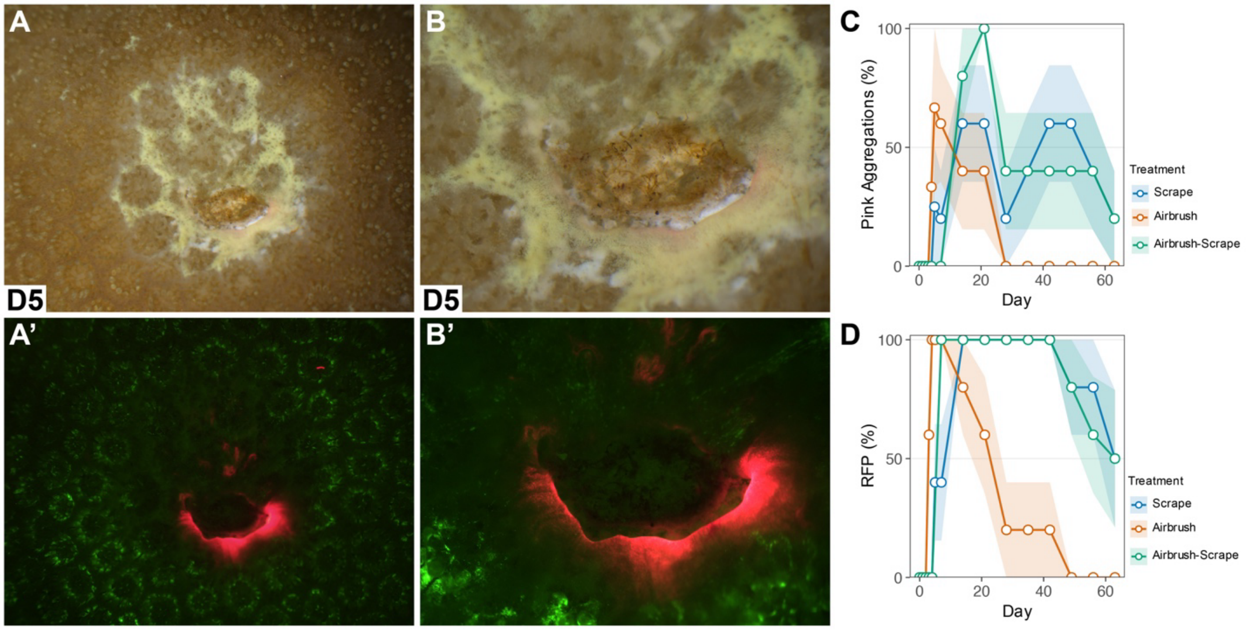

**Figure S2.** Wound-type differences in *Porites* emerge within the first few days, before the long-term trajectories diverge. Percentage of fragments showing each early-response feature over days 0–5, by wound type (mean across fragments; shaded bands ±1 SE; single-observation cells shown without a band). Scrape wounds begin accumulating coenosarc within roughly two days, whereas airbrush and airbrush-scrape wounds instead accumulate algal plugs over the same window — establishing the divergence in healing trajectory at the outset. Complements the main-text *Porites* time series (Figure 3).

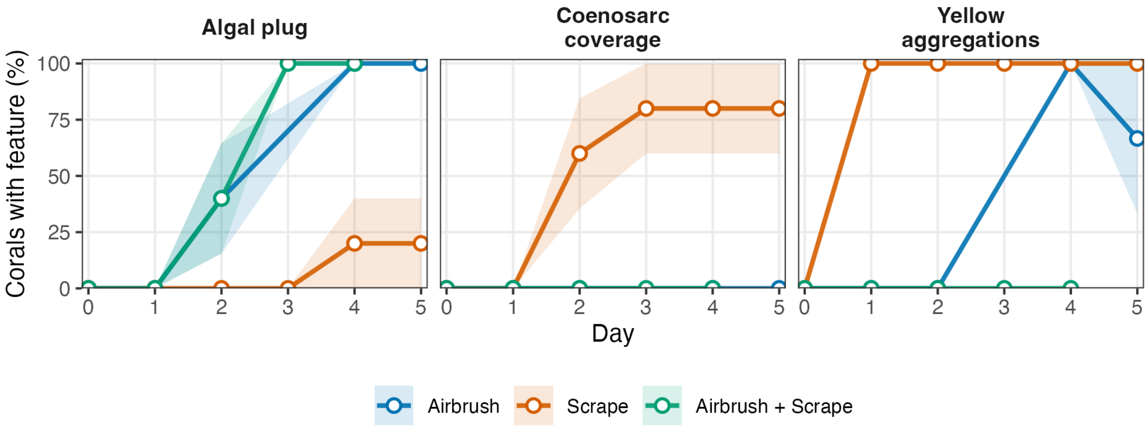

**Figure S3.** The reported Experiment 1 models fit the data adequately, with no sign of residual misspecification. Quantile–quantile uniformity (left) and residual-versus-predicted (right) plots from 1000 seeded DHARMa simulations for the healing, regeneration, and algal-colonization additive GLMMs. Residuals track the 1:1 line and show no uniformity, dispersion, or outlier departures (test statistics in Table S9), supporting the binomial GLMM specification used throughout. Cited in Methods (model adequacy).

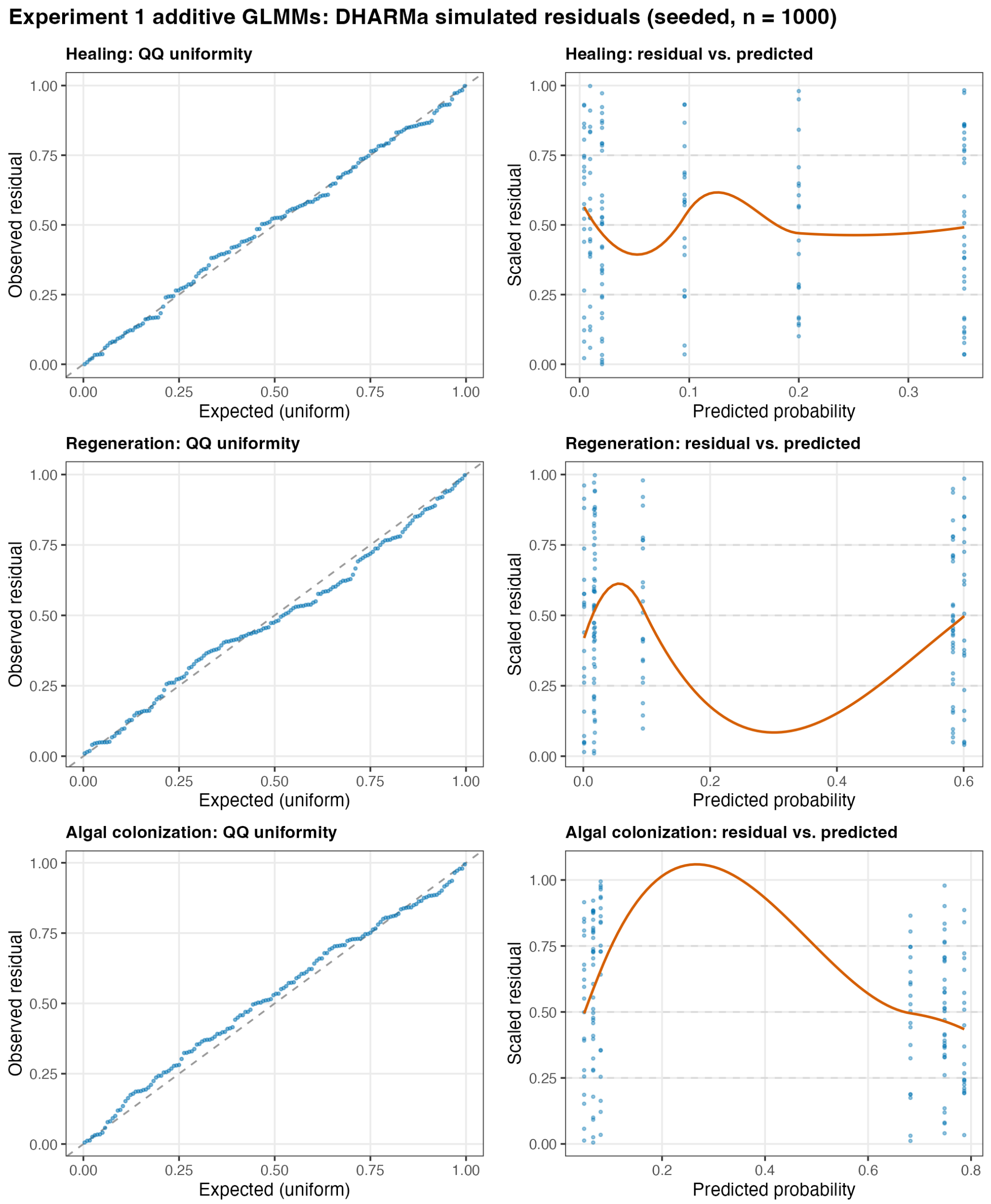

**Figure S4.** Residual diagnostics for the six Experiment 2 (*Porites*) time-series models, shown for completeness alongside the Experiment 1 diagnostics (Figure S3). Quantile–quantile uniformity (left) and residual-versus-predicted (right) plots from 1000 seeded DHARMa simulations for each scored outcome, using the separation-robust model adopted for inference (a penalized GLMM for every outcome except pink aggregations, a standard GLMM). Quantile residuals track the 1:1 line for every outcome, and the uniformity and outlier tests are non-significant throughout (all uniformity $P > 0.28$; all outlier $P = 1$); the dispersion test is significant for four of the six outcomes (coenosarc coverage, polyps, RFP, and yellow aggregations) but not for algal plug or pink aggregations. These simulation-based tests are low-powered and highly sensitive to sparsity in binary longitudinal data fit with weakly-informative priors, so they are reported descriptively rather than as a pass/fail gate — the treatment × time contrasts in Table S10 are unchanged across spline flexibility (Table S12). Cited in Experiment 2 Methods.

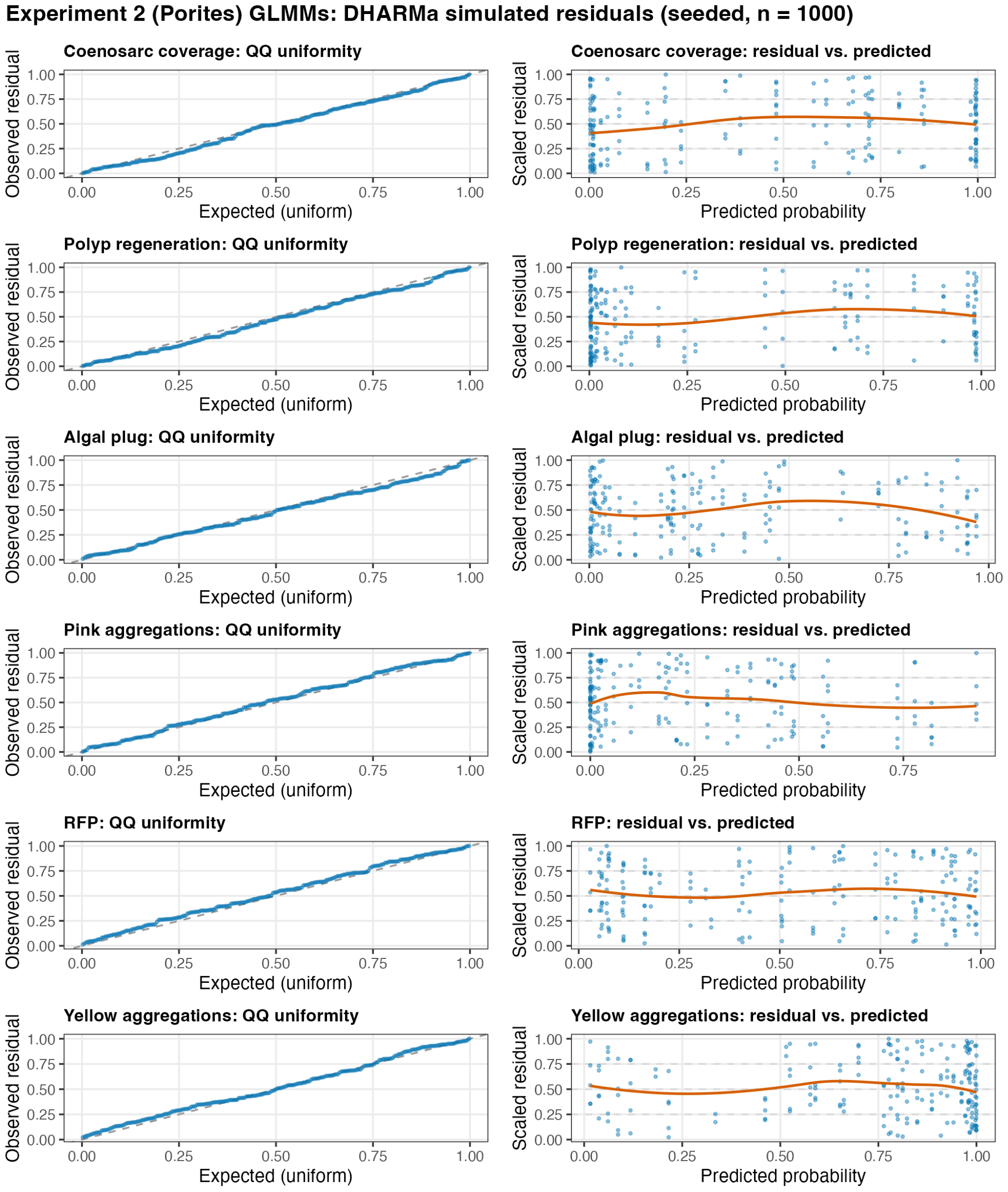
